# Plasticity of the proteasome-targeting signal Fat10 enhances substrate degradation

**DOI:** 10.1101/2022.07.15.499953

**Authors:** Hitendra Negi, Aravind Ravichandran, Pritha Dasgupta, Shridivya Reddy, Ranabir Das

**Author notes:** Electronic address. Currently at Institute for Stem Cell Science and Regenerative Medicine (inStem), Bangalore, India (Shridivya Reddy).

## Abstract

The proteasome controls levels of most cellular proteins, and its activity is regulated under stress, quiescence, and inflammation. However, factors determining the proteasomal degradation activity remain poorly understood. Proteasome substrates are conjugated with small proteins (tags) like ubiquitin and Fat10 to target them to the proteasome. It is unclear if the structural plasticity of proteasome-targeting tags influences substrate degradation. The tag Fat10 is activated during inflammation, and ambiguously, Fat10 and its substrates undergo rapid proteasomal degradation. We report that the rapid degradation of Fat10 substrates depends on its structural plasticity. While the ubiquitin tag is recycled at the proteasome, Fat10 is degraded with the substrate, and the mechanical unfolding kinetics of Fat10 regulates substrate degradation. Our studies reveal that long-range salt bridges are absent in the Fat10 structure, creating a plastic protein with partially unstructured regions suitable for proteasome engagement. Such a malleable structure also provides low resistance to mechanical unfolding and expedites proteasomal degradation. We also uncovered that the Fat10 plasticity destabilizes substrates significantly and creates partially unstructured regions in the substrate to enhance degradation. NMR-relaxation-derived order parameters and temperature dependence of chemical shifts identify the Fat10-induced partially unstructured regions in the substrate. They correlated excellently to the regions where Fat10 contacts the substrate, suggesting that the tag-substrate collision destabilizes the substrate. These results highlight a strong dependence of proteasomal degradation on the structural plasticity and thermodynamic properties of the proteasome-targeting tag.

## INTRODUCTION

Proteasome substrates must be conjugated with other small proteins, known as proteasome-targeting tags, to target substrates to the proteasome ^1^. Ubiquitin is a proteasome-targeting tag conjugated to substrates by posttranslational modification. It interacts with the 19S proteasome subunit, which harbors ubiquitin receptors. Consequently, 19S aligns its base subunits, consisting of AAA+ ATPases, with the 20S Core Particle (CP) to create a channel for substrate entry. Deubiquitinase enzymes cleave the ubiquitin, while the substrate enters the ATPases to unfold and translocate to CP, where it is cleaved into short peptides by proteases ^2–4^. Proteasomal degradation regulates the levels of most nuclear and cytosolic proteins^5^. Global cellular protein degradation rate is modulated in the quiescent state, under nutrient stress, or during inflammation^5–8^. However, mechanisms that regulate proteasome degradation activity in such conditions remain poorly understood. Partially unstructured regions at the N/C-termini or within the substrate are critical to proteasome engagement and degradation. Global disorder and topology of substrates also influence the degradation rate^9^. It is unclear if the plasticity of the proteasome-targeting tag can regulate proteasome activity and substrate degradation rate.

During stress or inflammation, a swift change in proteasomal degradation activity by changing the global disorder in proteins or its biased sequences may be challenging. An alternate mechanism is to upregulate a proteasome-targeting tag that rapidly degrades substrate proteins. The human leukocyte antigen-F adjacent transcript 10 (Fat10) is a proteasome-targeting tag that directly targets substrates for proteasomal degradation^10^. Fat10 expression is restricted to immune cells^11^, and other cell types express Fat10 upon induction by proinflammatory cytokines^12–14^. Fat10 includes two ubiquitin-like domains attached by a flexible linker and is conjugated to substrate proteins by posttranslational modification. Fat10’s surface properties and interactions are distinct from ubiquitin^15^. Fat10 has a lower half-life of 1 hr, while ubiquitin has a longer half-life (∼24 hrs)^16,17^. Fat10’s shorter half-life prevents a prolonged inflammatory response that can potentially lead to apoptosis^17^. Mechanisms underlying the rapid degradation of Fat10 are poorly understood.

The substrates of Fat10 overlap significantly with ubiquitin substrates^17^, suggesting that Fat10 is an auxiliary proteasomal-targeting signal activated during inflammation. However, several mechanistic differences exist between the ubiquitin- and Fat10-proteasome pathways. Ubiquitin is cleaved by deubiquitinating enzymes at the proteasome before the substrate enters ATPases, but Fat10 remains uncleaved and is degraded along with the substrate^13^. Well-folded substrates must be globally or partially unfolded for engagement with the proteasome. Accessory unfoldases like Cdc48 assist the process by unfolding substrates^18^ before they interact with the proteasome. Substrate unfolding by accessory unfoldases is necessary for the ubiquitin substrates but not for the Fat10 substrates^15^, suggesting a profound impact of Fat10 on the substrate’s structure and energetics. However, Fat10’s effect on the substrate’s structure and thermodynamics is unknown.

While ubiquitin has a rigid structure, Fat10 has a flexible fold, providing an opportunity to probe if the plasticity of proteasome-targeting tags affects substrate degradation rate. In this work, we correlate the thermodynamic properties and conformational plasticity of Fat10 and ubiquitin with the proteasomal degradation rate of their substrates. Our results suggest that the Fat10 free energy barrier is substantially lower than ubiquitin. Consequently, the Fat10 unfolding kinetics is seven-fold faster than ubiquitin, and Fat10’s mechanical resistance to unfolding is weaker than ubiquitin. The absence of long-range salt bridges in Fat10 creates partially unstructured regions, leading to efficient proteasome engagement and degradation. Furthermore, Fat10’s structural plasticity reduces the thermodynamic stability of substrate proteins in cellular and *in-vitro* conditions and creates local partially unstructured regions in the substrate. The substrate reciprocally reduces Fat10 stability by thermodynamic coupling. These destabilization effects in Fat10 and the substrate work synchronously to create more partially unfolded regions in the conjugate and enhance degradation. NMR experiments measured enthalpy and conformational entropy to reveal Fat10-induced sites of local disorder in the substrate. Experimental and computational studies suggest that nonspecific collisions with the proteasome-targeting tag destabilize the substrate. These results provide the underlying mechanism of the rapid degradation of Fat10 substrates and highlight that the proteasome-targeting tag’s conformational plasticity regulates proteasomal degradation.

## Materials and Methods

### Unfolding studies

Chemical denaturation studies by CD spectroscopy were carried out using 20 µM of protein (Ub, SUMO1, Fat10, Fat10D1, and Fat10D2) incubated overnight with various concentrations of Guanidium Chloride (GdnCl, 0 to 6M) made in the native buffer at 25 °C. The change in the far-UV signal at 222nm was monitored using the Jasco J-1500 spectropolarimeter. CFP and its variants were studied similarly by incubating 15 nM of protein with GdnCl. CFP fluorescence was measured using Horiba Fluromax-4 with an excitation wavelength of 434 nm. The emission signal was collected at 474 nm. Ub_F45W_ and its variants were similarly incubated with GdnCl. The signal at 340 nm was monitored for the change in intrinsic tryptophan fluorescence. Chemical denaturation data were normalized to a two-state unfolding equation using SIGMA plot software. The final curve fitting used a monomeric two-parameter melt equation (Privalov, 1979). The calculated free energy of unfolding, ΔG_unfolding_, and slope of the transition curve, m, were used to calculate free energy (ΔG_0_) at 0M [GdnCl] using the equation ΔG_unfolding, [GdnCl]_ = ΔG_0_ – m[GdnCl].

Proteins were dialyzed to phosphate buffer (50 mM, pH 6.5, 250 mM NaCl) with or without DTT (1 mM) for thermal equilibrium unfolding studies. The unfolding transition was monitored by observing the change in the CD signal at 222nm on the JASCO J-1500 spectropolarimeter connected to a Peltier. Data were collected from 20°C to 95°C after every 1°C per minute rise in temperature with 32 sec of data integration. Raw data of the CD signal were normalized to a two-state unfolding equation and plotted against temperature. The curve was fitted with a sigmoidal equation to yield T_m_. Thermal equilibrium unfolding data were analyzed and processed using SIGMA plot software.

### Native-state Proteolysis

The protocol for native-state proteolysis has been described previously^19^. 100 μl aliquots of Ub-CFP and Fat10-CFP were treated with multiple thermolysin protease concentrations (stock concentration 10 mg ml^-1^). At different time points, 10 μl of the reaction mixture was taken, quenched with 6x-SDS loading dye, and run on SDS-PAGE gels. An I-Bright imaging instrument captured the in-gel fluorescence image (Life Technologies). Images were inverted to greyscale and quantified by normalizing them to no protease control sample. The quantified data were fitted to the first-order exponential equation and observed proteolysis kinetics (k_obs_) were calculated at different thermolysin concentrations. The mean of k_obs_ (n=3) at different concentrations was fitted to a linear equation against thermolysin concentration. ΔG_proteolysis_ was calculated using Eq. 3 by using the slope of the linear fit (Eq. 2). The k_cat_/K_M_ of Thermolysin is 99,000M^-1^ s^-1^ (Park and Marqusee, 2004).

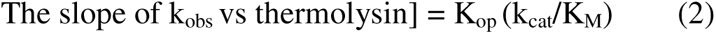

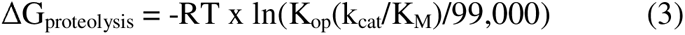

### NMR studies

^13^C,^15^N isotopically labeled Fat10D1, and Fat10D2 were prepared in 50 mM Tris, 250 mM NaCl buffer with pH 7.5. Standard triple resonance multidimensional NMR experiments ^15^N-HSQC, HNCA, HN(CO)CA, CBCACONH, and HNCACB were recorded at 298K on an 800MHz Bruker Avance III HD spectrometer equipped with a cryoprobe head. These experiments yielded the backbone assignment of Fat10D1 and Fat10D2. For proteins Fat10D1-Ub and Ub-Fat10D2, only the ubiquitin residue peaks were assigned using standard triple resonance experiments HNCA, HNCOCA, CBCACONH, and HNCACB. NMR data were processed in NMRpipe^20^ and analyzed by NMRFAM-SPARKY^21^ software. After peak-picking of the backbone experimental data in SPARKY, the peaks were assigned by the PINE server^22^ and confirmed manually.

### Cell Culture Experiments

Approximately 0.1 million cells/well of a 12-well plate were seeded with HEK293T cell line and transfected after 70% confluency using a Lipofectamine reagent (Promega). For the cycloheximide assay, cells were transfected with 3xFLAG-wtFat10-AV, 3xFLAG-wtFat10D1-CYCIAV, 3xFLAG-wtFat10D2-AV, and 3xFLAG-UbK0GV. The transfected cells were incubated at 37°C for 18-20 hours, treated with Cycloheximide (CHX, final concentration 50 µg/mL), and lysed at different time points. For six hours, these cells were treated with MG132 (final concentration 10 µM). 20 mM Tris (pH 7.5), 150 mM NaCl, 1X protease inhibitor cocktail, and 1% NP40 were used to lyse the cell pellet. The BCA kit (Thermo) estimated the total protein amount. Similar transfections were carried out for FLAG-UbGV-GFP and FLAG-Fat10AV-GFP. After 18hrs of incubation, the cells were treated with CHX (100 μg/mL) in the presence and absence of p97 inhibitor (DBeQ; final concentration 10µM) and further incubated for different time points. All the Immunoblots were probed with Mouse anti-FLAG antibody (SIGMA, 1 in 10,000 dilutions) and Mouse anti-β-actin antibody (Santa Cruz, 1 in 5000 dilutions) and further probed with HRP conjugated anti-mouse secondary antibody at 1:10,000 dilutions. The blots were developed using clarity ECL (Bio-Rad) staining reagent, and the chemiluminescence signal was observed in Image Quant LAS 4000 (GE). The Immunoblots for GFP variants were probed similarly with the mouse anti-FLAG antibody. Tubulin was used as the loading control for normalization and probed with Mouse anti-Tubulin antibody (SIGMA, 1 in 3000 dilutions). All the immunoblots of GFP variants were developed, and the chemiluminescence signal was observed in Amersham Imager 400 (GE). Quantification was done using ImageJ software with three biological replicates for each experiment.

### In-cell protein stability assay

The stability of CRABP1 with ubiquitin and Fat10 in the cellular environment was monitored by pre-labeling the *E. coli* BL21 (DE3) cells with FIAsH-EDT_2_ fluorescent dye (final concentration 150µM) at OD_600_ = 0.5, followed by incubation till OD_600_ reaches 1. IPTG was added to cultures to induce protein synthesis. After two hours, the culture solution was aliquoted (150 µl) and treated with various concentrations of Urea ranging from 0M to 3M, keeping the final volume at 400 µl. The culture solutions were incubated for 30 mins before pelleting and washing with native buffer (10 mM Tris buffer, pH 7.5), followed by the fluorescent measurements (excitation 500 nm and emission 531 nm). The rest of the protocol is the same as described previously (Ignatova and Gierasch, 2004).

### High-temperature unfolding simulations

Systems were prepared in a cubic box of TIP3p water, with a minimum distance of at least 50 Å between solute atoms and the box edge. Counter ions were added to neutralize the system. The system setup and equilibration procedures were similar to the unrestrained simulations, with 450 K as the equilibration temperature. Protein unfolding simulations were performed at 450 K for 300 ns. The simulations were performed with ten replicas to obtain an average folding pathway.

### ASMD simulations

Constant force explicit solvent ASMD simulations of Fat10D1, Fat10D2, Ub, full-length Fat10, and Ub_2_ were carried out. The reaction coordinate is the end-to-end distance between the Cα atoms of the first amino acid and the last amino acid of the respective protein. For simulations consisting of Fat10D1, Fat10D2, and Ub, the proteins were pulled at 1 Å/ns velocity, and for full-length Fat10 and Ub_2_, the pulling velocity was 5 Å/ns. The reaction coordinate was partitioned into ten equal segments, each with ten trajectories. The system was energy-minimized equilibrated, and the resulting coordinates & velocities were used as starting points for ASMD simulations. The simulations were performed with periodic boundary conditions in the NPT ensemble; electrostatic interactions were computed by the PME (Particle Mesh Ewald) method. Nonbonded interactions were treated with a cutoff of 8 Å.

Further details of molecular biology, protein purification, MD simulations, and biophysical experiments are provided in the Supporting Information.

## Results

### Fat10 undergoes rapid proteasomal degradation

Although UBLs are structurally similar, they are diverse in sequence landscape. The N-terminal and C-terminal Fat10 domains (Fat10D1 and Fat10D2) share 29% and 36% sequence identity with ubiquitin, respectively (Figure 1A-C). Moreover, the two domains of Fat10 share only 18% sequence identity between themselves. Important regions like the L8-I44-V70 hydrophobic patch, an interaction hotspot in ubiquitin, are absent in Fat10. A phylogenetic tree analysis based on available ULM structures showed that among the two Fat10 domains, the C-terminal Fat10D2 is structurally closer to ubiquitin (Figure S1A-S1D). Given the poor sequence identity between ubiquitin and Fat10, their cellular levels could be distinctly regulated.

**Figure 1.**
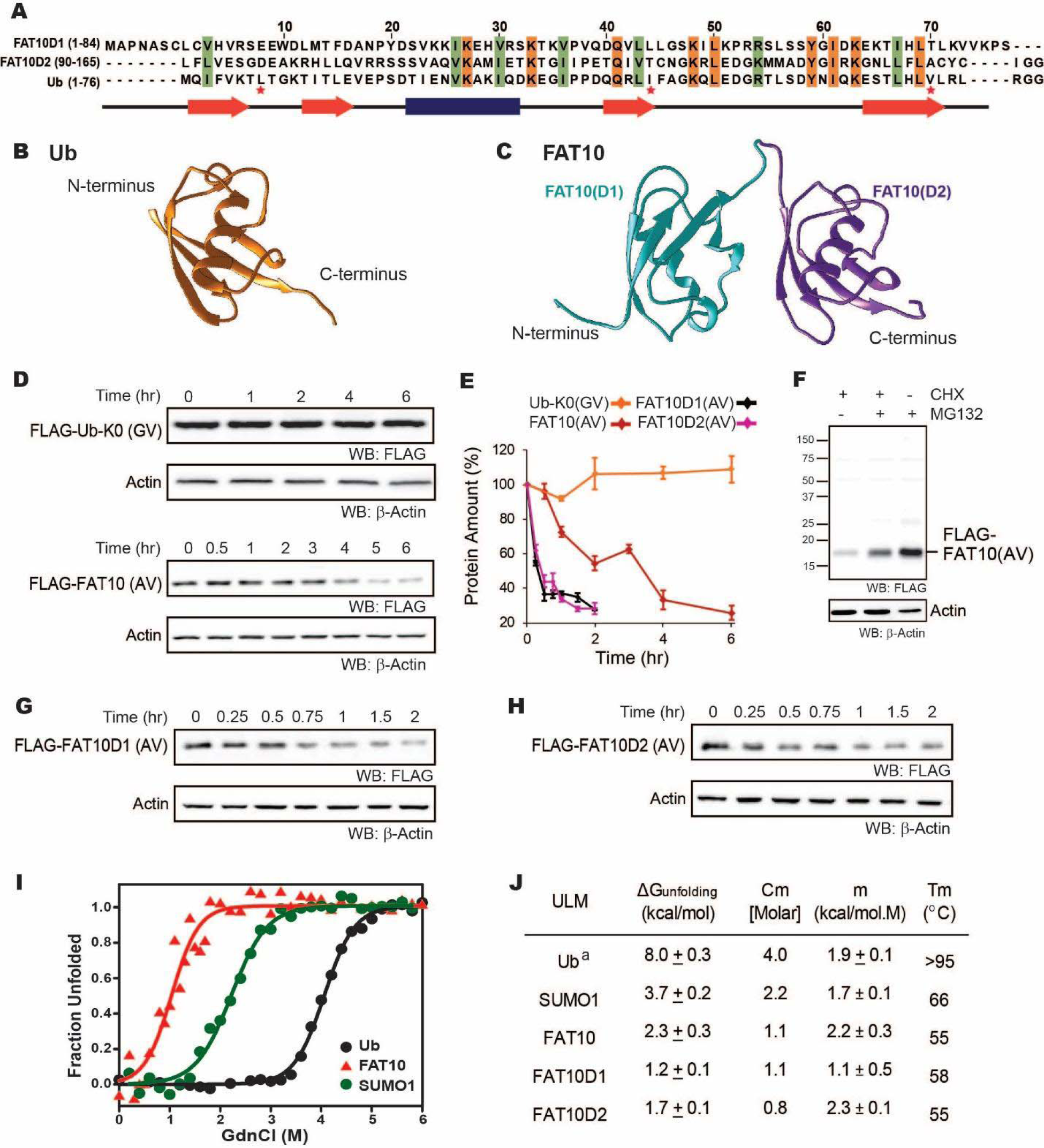
(A) Structure-based sequence alignment of Fat10D1 (PDB: 1gf1), Fat10D2 (PDB: 1gf2), and ubiquitin (PDB: 1ubq). Conserved hydrophobic and identical residues in Fat10D1, Fat10D2, and ubiquitin are highlighted in light green and orange colors. The L8-I44-V70 residues that create a ‘hot spot’ of interactions in ubiquitin are marked with a red asterisk. (B) Structure of ubiquitin (1UBQ; orange) and (C) Homology model structure of full-length Fat10 where Fat10D1 is colored as cyan, and Fat10D2 is colored as purple. (D) FLAG-UbK0(GV) and FLAG-Fat10(AV) protein levels are plotted against time. The C-terminal GG residues are substituted with GV or AV to prevent conjugation to the cellular substrates. HEK293T cells were transfected with either FLAG-Ub or FLAG-Fat10(AV), treated with Cycloheximide, and lysed at different time points. The lysates were separated on SDS PAGE gels and blotted with anti-FLAG antibodies. (E) Quantified protein levels in (D) are plotted against time after normalizing with β-Actin. (F) HEK293T cells were transfected with Fat10, treated with/without Cycloheximide, and proteasomal inhibitor MG132. The lysates were separated on SDS PAGE gels and blotted with anti-FLAG antibodies. (G) Similar to (D), showing degradation of FLAG-Fat10D1(AV) & (H) FLAG-Fat10D2(AV) after cycloheximide treatment. Quantified protein levels of FLAG-Fat10D1(AV) and FLAG-Fat10D2(AV) are plotted in (E). (I) GdnCl melt curves of Fat10, Ub, and SUMO1. Normalized mean ellipticity shift is plotted against GdnCl concentration. (J) A table with the stability parameters of Ub, SUMO1, Fat10, and Fat10 domains is provided. (a: Reference ^23^).

The degradation rate of ubiquitin and Fat10 was compared in cells using cycloheximide chase assay. Non-conjugable variants UbK0-GV and Fat10-AV were expressed in HEK293T3 cells, and the protein levels were measured against time (Figure 1D). Fat10 levels dropped sharply compared to ubiquitin, suggesting that Fat10 undergoes rapid degradation (Figure 1E). Treatment with a proteasomal inhibitor restores Fat10 (Figure 1F), indicating that Fat10 undergoes proteasomal degradation. The protein levels of individual Fat10 domains were also studied. Both domains degraded faster than the full-length Fat10 (Figure 1E, G, H), suggesting that the Fat10 is more stable than the individual domains. The isolated Fat10 domains were also degraded via the proteasomal pathway (Figure S1E-F). Protein cofactors specific to the full-length Fat10 may bind and bury its partially disordered regions to reduce proteasomal degradation. Transient interactions between the isolated domains may also bury the disordered region and provide a more compact structure to the full-length Fat10, reducing its degradation.

### Thermodynamic characterization of Fat10

To study the correlation between Fat10 degradation and its structural and thermodynamic properties, Fat10 unfolding was studied in a purified system. Fat10 was recombinantly expressed and purified. The far-UV Circular Dichroism (CD) spectra and two-dimensional ^15^N Heteronuclear Single Quantum Coherence (^15^N-HSQC) NMR spectra of Fat10 suggested a well-folded tertiary structure (Figure S2A, B). The thermodynamics of unfolding Fat10 and other ULMs were measured by chemical denaturation using guanidinium hydrochloride (GdnHCl). Gibbs free energy of unfolding (ΔG_unfolding_) for ubiquitin was 8 kcal/mol (Figure 1I-J). ΔG_unfolding_ for another ULM, SUMO1 was lower than ubiquitin (3.7 kcal/mol). However, ΔG_unfolding_ was lowest for Fat10 (2.3 kcal/mol, Figure 1I, 1J). Ubiquitin also has high thermodynamic stability, and its melting temperature (Tm) is greater than 95°C ^23^, whereas Fat10 unfolded with a Tm of 55°C (Figure S2C), suggesting Fat10 has a malleable structure compared to ubiquitin. Differential scanning fluorimetry measurements had reported similar differences in Tm between Fat10 and ubiquitin^15^.

Transient interactions between the Fat10 domains may bury partially disordered regions and increase the thermodynamic stability of the full-length protein. In that case, the unfolding energies of individual domains shall be lower than the full-length protein ^24^. The two Fat10 domains were isolated, and their energetics was measured. Far-UV CD and ^15^N-HSQC spectra confirmed that the purified isolated Fat10 domains are folded (Figure S3A-S3C). GdnHCl-induced and temperature-induced denaturation of the Fat10 domains were carried out (Figure S3D-S3F). ΔG_unfolding_ of individual domains is lower than full-length Fat10 (Figure 1J), suggesting transient interactions between the Fat10 domains stabilize the full-length protein. ΔG_unfolding_ and Tm values indicate that Fat10 has substantially lower thermodynamic stability than ubiquitin.

### Fat10 has low resistance to mechanical unfolding

Protein mechanical unfolding by molecular motors, such as the proteasomal ATPases, can be simulated by steered molecular dynamics^25^. The mechanical unfolding of Fat10 was studied in the explicit solvent by Adaptive Steered Molecular Dynamics (ASMD) (Figure 2A, Movie S1, Figure S4)^26^. A linear Di-ubiquitin (Ub2) was used for comparison, whose size is similar to Fat10. While the work required to unfold and linearize Ub_2_ was 735 kcal/mol, the same for Fat10 was 581 kcal/mol (Figure 2B). The lower work required to unfold Fat10 corroborated the lower value of ΔG_unfolding_ measured during Fat10 denaturation (Figure 1J). The simulation was repeated for each isolated Fat10 domain and compared to monoubiquitin. The work required to unfold the isolated Fat10 domains was lower than ubiquitin by 30-60 kcal/mol (Figure 2B, Figure S5). Resistance during mechanical unfolding arises from cooperative packing interactions between buried sidechain atoms^27^. The lower resistance in Fat10 suggests weaker cooperative packing interactions at the protein’s buried core.

**Figure 2.**
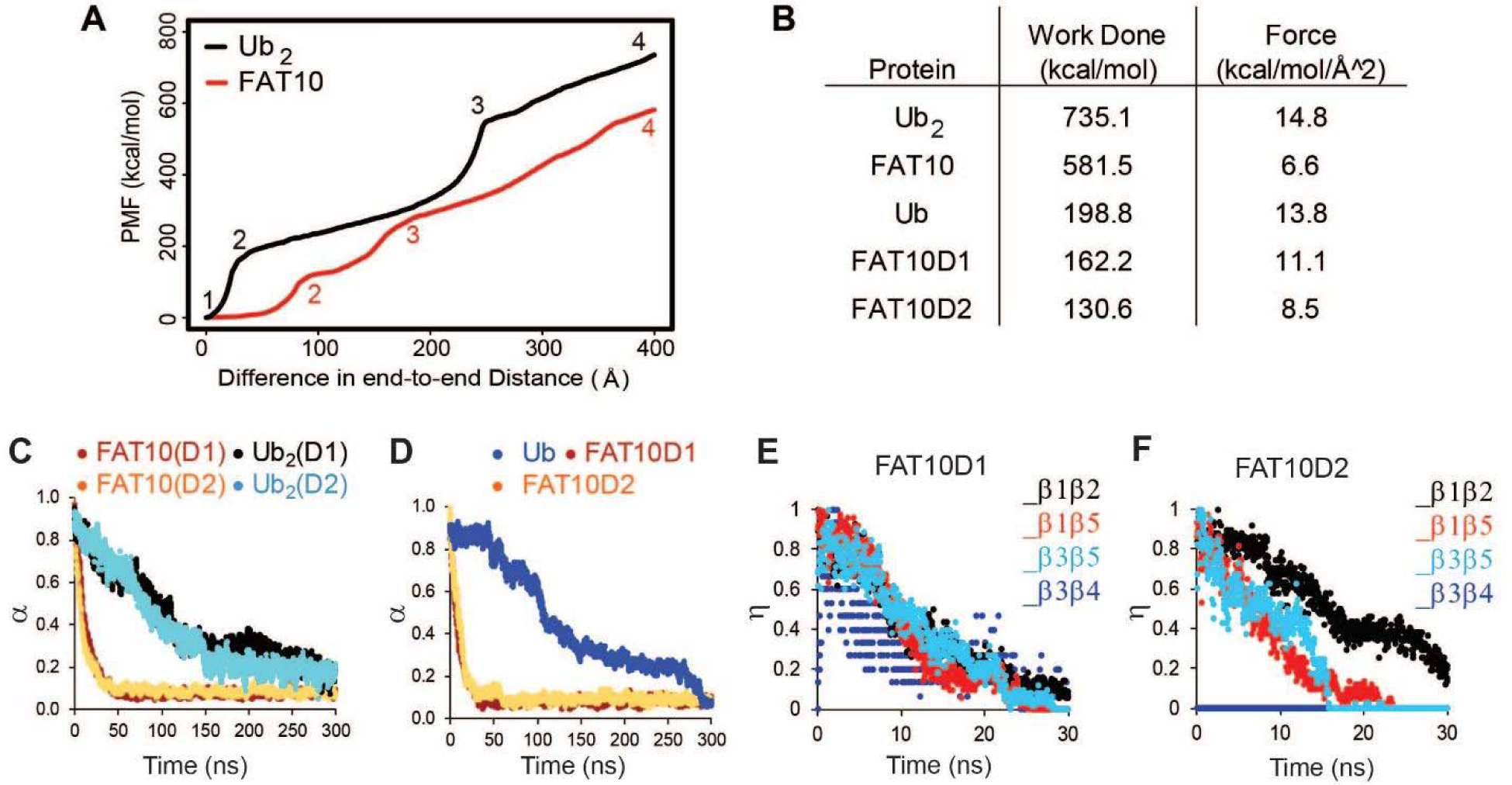
Unfolding studies of Fat10, di-ubiquitin (Ub_2_), Ub, and individual domains D1/D2 in Fat10 by MD simulations. (A) ASMD of Ub_2_ and Fat10. The potential Mean Force (PMF) is plotted against the normalized end-to-end distance. Each unfolding event is marked by a number, whose corresponding conformation is given in Figure S4. (B) The work done to unfold the proteins by ASMD is provided. (C) Simulations of Fat10, di-ubiquitin (Ub_2_), Ub, and individual domains D1/D2 in Fat10 were performed at 450K. The fraction of native contacts is defined as (α) and plotted against time. Fat10(D1) and Fat10(D2) are the D1 and D2 domains in Fat10. Ub_2_(D1) and Ub_2_(D2) are the two Ub domains in diubiquitin. The data is averaged over ten replicas. (D) is the same as (C) measured for individual domains Fat10D1, Fat10D2, and Ub. (E) The fraction of native pairwise backbone hydrogen bond is defined as (η), plotted for β1 to β5 of Fat10D1. (F) The same as (E) for the Fat10D2 domain.

Weak cooperative intramolecular interactions should result in faster unfolding kinetics. Fat10 unfolding kinetics was studied by MD simulations at high temperatures. 4 μs simulations were performed at 450K for Ub_2_ and Fat10 (Movie S2). Native intramolecular contacts, including Van der Waals (VdW) interactions and hydrogen bonds (hbonds), decrease over time as the protein unfolds at high temperatures. While Ub_2_ unfolded gradually in 300 ns, Fat10 unfolded drastically within 40 ns (Figure 2C). The ULM proteins adopt a β-grasp fold consisting of a five-strand β-sheet. The β-sheet hbonds were disrupted promptly in Fat10 as opposed to ubiquitin (Figure S6A). The native contacts and hbonds in the individual isolated Fat10 domains also disrupted faster than ubiquitin (Figure 2D and S6B), confirming weak cooperative interactions in Fat10.

The fraction of native β-sheet hbonds in Fat10 and ubiquitin were plotted in Figures 2E, 2F, and S6C. Distinct sequences of inter β-strand hbonds disruption between the Fat10 domains and ubiquitin suggest distinct unfolding pathways. A comprehensive analysis of the secondary structure hbonds during unfolding showed that the 3^10^ helix α2 unfolds first in ubiquitin, followed by β3-β4, α1, β3-β5 and β1-β5 contacts (Figure S6D). The β1-β2 contacts break at the last step. The unfolding pathway is similar to that observed in equilibrium simulations of ubiquitin^28^. In contrast, the first unfolding event in Fat10D1 is the disruption of β3-β4 contacts, followed by α2, α1, and β1-β5 contacts (Figure S6E). The β3-β5 and β1-β2 contacts are disrupted simultaneously at the last step. Similarly, the unfolding pathway of the second domain in Fat10 is distinct from ubiquitin (Figure S6F). Altogether, the resistance to mechanical unfolding is lower in Fat10 than in ubiquitin, the kinetics of Fat10 unfolding is faster, and the Fat10 unfolding pathway is distinct from the ubiquitin unfolding pathway.

### The absence of key interactions creates partially unstructured regions in Fat10

The flexible Fat10 structure may sample higher energy states with partially unstructured regions, which was investigated by MD simulations at room temperature. Root Mean Square Fluctuations (RMSF) of Fat10 domains are high at the β1-β2 loop, the C-terminal end of α1, and the α2 loop, indicating regions with high flexibility (Figure S7). 2D free energy of Fat10 conformations was calculated as a function of the negative logarithm of root mean square deviation (RMSD) and the radius of gyration (Rgyr) populations. Ubiquitin was used as a control in these simulations. The 2D free energy plot of ubiquitin reflected a single highly populated state (Figure 3A), whose representative structure superimposes with the native ubiquitin x-ray structure (Figure 3B). In contrast, the native states of Fat10 domains are in equilibrium with higher energy partially unfolded states (Figure 3A). Local interactions between β1-β2 and helix α1 are disrupted, and loops at the N- and C-termini of α1 are disordered in these partially unfolded states.

**Figure 3.**
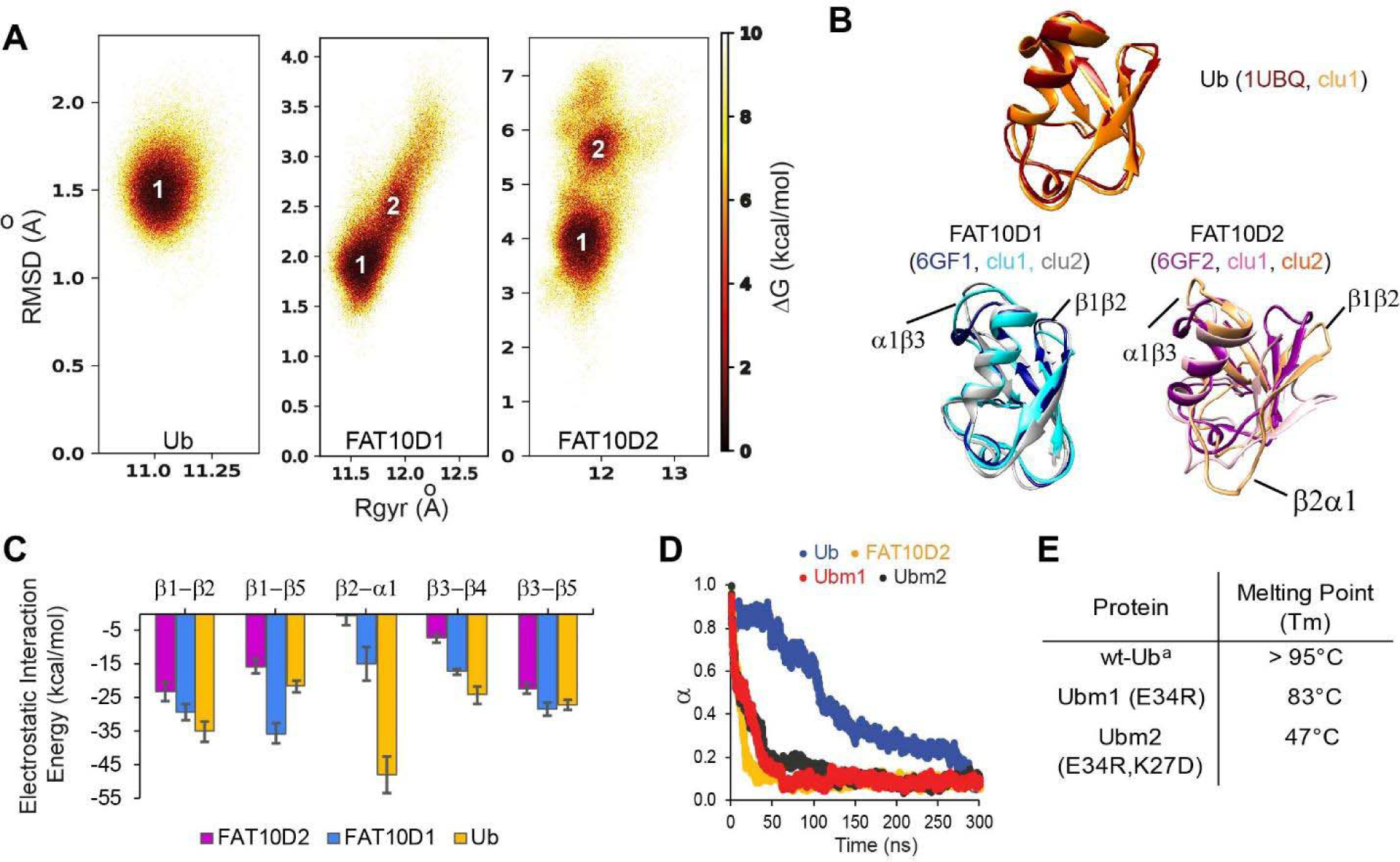
(A) The free energy landscape of Fat10 domains and ubiquitin are plotted as a function of RMSD and radius of gyration (Rgyr) obtained from simulations across three replicas (3 x 2.5 μs) performed at 300K. The minima from each cluster are numbered. (B) The corresponding conformation of each cluster in (A) is shown. The structures of the proteins, denoted by their pdb ids, are provided for comparison. (C) The mean electrostatic energy of interactions between different pairs of secondary structures obtained from simulations is plotted for Fat10 domains and ubiquitin. The error bars denote the standard deviation. (D) The fraction of native contacts in Ubm1, Ubm2, Ub, and Fat10 domain against time at 450K simulations. (E) The melting point of ubiquitin mutants. (a: Reference ^23^).

The radius of gyration (Rog) reflects the nature and extent of packing interactions in the protein. Although ubiquitin and the Fat10 domains are similar in size, the Rog of the Fat10 ground state is higher than ubiquitin, suggesting more flexibility and lower packing (Figure S7C). VdW and electrostatic interaction energies were calculated between the interacting secondary structures in Fat10 and ubiquitin to study the basis of reduced packing interactions. The VdW interaction energy was comparable between Fat10 domains and ubiquitin (Figure S7D). The electrostatic interaction energy was also similar in all regions except between β2 and α1, where Fat10 has a significantly lower negative value than ubiquitin, suggesting fewer electrostatic contacts (Figure 3C). Three hydrogen bonds and a salt bridge between β2 and α1 are present in ubiquitin but absent in Fat10, which reduces the interaction between β2 and α1 (Figure S8A). Another salt bridge between the α2 loop and α1 is also exclusive to ubiquitin (Figure S8B). These salt bridges may be crucial to the stability and compactness of ubiquitin, and their absence makes Fat10 flexible.

When these salt bridges were disrupted in ubiquitin by substitution, the free energy landscape changed, giving rise to partially unfolded forms similar to Fat10 (Figure S9A). The RMSF values of mutant ubiquitin increase at the C-terminal end of α1 and the β4-α2 loop (Figure S9B, S9C). Principle Component Analysis (PCA) and the RMSF values suggested that the ubiquitin mutants have increased dynamics in the regions of broken salt bridges (Figure S9D-S9F). When the unfolding kinetics were compared between mutants and wt protein by simulations at 450K, the mutants unfolded much faster than ubiquitin but similar to Fat10 domains (Figure 3D). The thermodynamic stability of ubiquitin mutants was measured experimentally. Disrupting the salt bridges reduces the melting point of ubiquitin by 50°C, suggesting the importance of the salt bridges for ubiquitin stability (Figure 3E and Figure S9G). The salt bridges were engineered in the N-terminal Fat10 domain by appropriate substitutions. The protein with engineered salt bridges populated the ground state but not the partially unfolded states (Figure S10). RMSF and PCA analysis suggested that engineered salt bridges reduced the fluctuations and increased rigidity in the domain (Figure S10). Overall, critical electrostatic interactions present between α1 and β1β2 and between α1 and α2 loop stabilize the ubiquitin fold. These interactions are absent in Fat10, creating a flexible structure that is in dynamic equilibrium with partially unfolded forms.

### Weak hydrogen bonds and high conformational entropy detected in Fat10

Protein thermodynamic stability depends on the energy of enthalpic interactions and conformational entropy. The temperature dependence of backbone amide proton chemical shifts is an excellent reporter of backbone hbond strength, which contributes to enthalpic interactions.^29^. The temperature dependence of the amide proton chemical shifts is the temperature coefficient Tc, where Tc = Δδ^NH^/ΔT ^30^. Higher negative Tc values suggest weaker hbonds and disorder propensity, while lower negative Tc values suggest stronger hbonds and structural rigidity. Tc’s were measured in Fat10 and plotted against each residue in Figure 4A, and the fitting errors are provided in Figure S11A. In the N-terminal Fat10 domain, residues in the region β1 to β2 and the loop between including α2 had high negative Tc values, suggesting weaker hbonds and disorder propensity in these regions (Figure 4A). Similarly, several residues in the β1, β2, α1, and α2 loop in the C-terminal domain had high negative Tc values.

**Figure 4.**
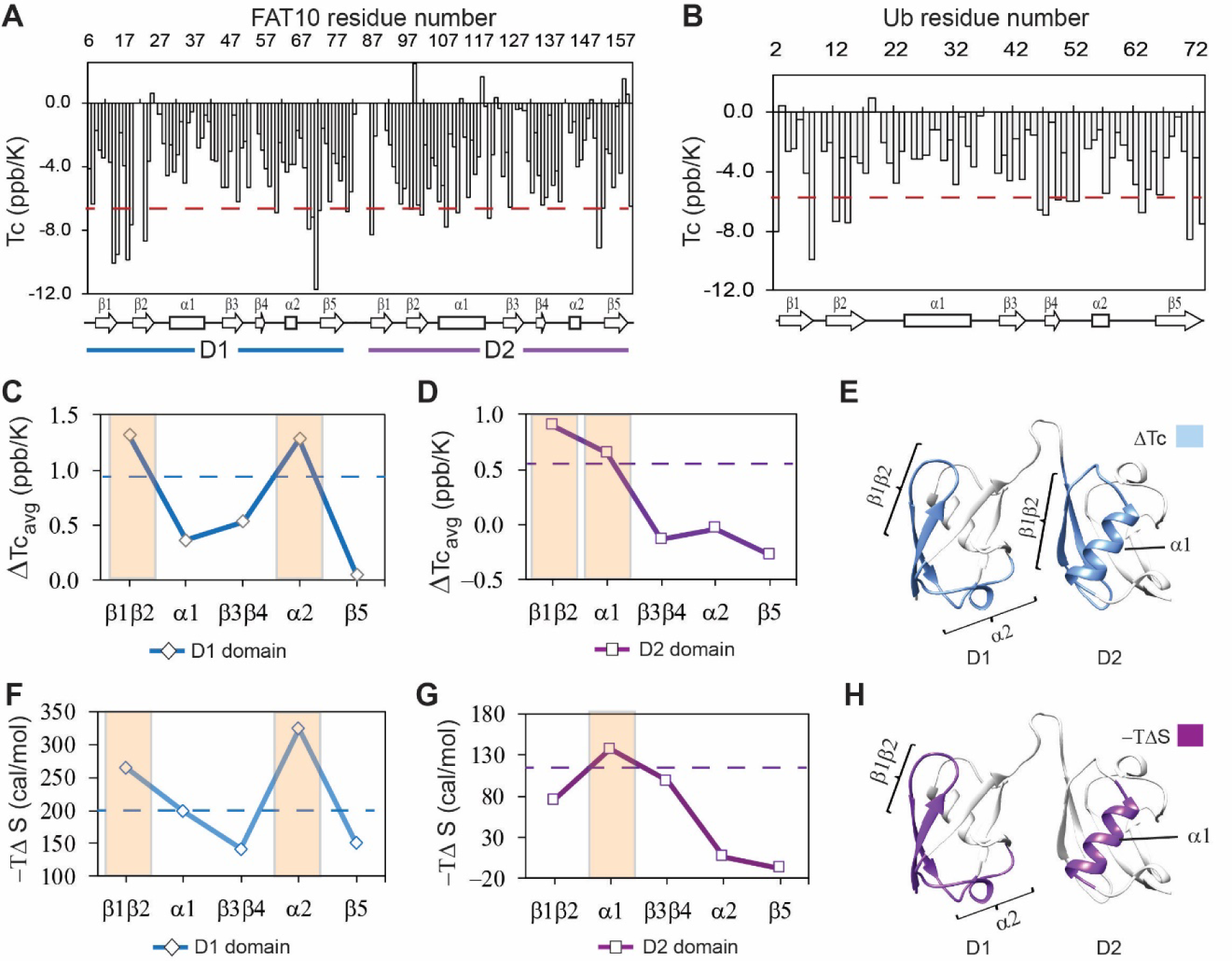
The local hbond stability and conformational entropy in Fat10. Temperature coefficients (Tc) are plotted for Fat10 and ubiquitin in (A) and (B), respectively. The horizontal red line is (mean – S.D.), where the mean value is negative. High negative Tc values suggest weaker hbonds and disorder propensity. (C) The difference in average temperature coefficients (ΔTc_avg_) between the N-terminal Fat10 domain (D1) and ubiquitin, where ΔTc_avg_ = Tc(Ub)_avg_ - Tc(D1)_avg_. The blue dashed line is mean + error. (D) The difference in averaged temperature coefficients (ΔTc_avg_) between the C-terminal domain and Ub, where ΔTc_avg_ = Tc(Ub)_avg_ - Tc(D2)_avg_. The purple dashed line is the mean + error. Higher ΔTc_avg_ values suggest weaker hbonds and destabilization in these Fat10 regions than ubiquitin. (E) The segments with high ΔTc_avg_ values are colored light blue on the Fat10 structure. (F) The difference in conformational entropy -TΔS_conf_, where ΔS_conf_ = S_conf_^Ub^ – S_conf_^D1^, was averaged for the various segments and plotted. The entropy values were calculated from the order parameters measured in Figure S12. The broken line denotes (mean + error). (G) Same as (F) except conformational entropy is calculated for the C-terminal Fat10 domain, such that ΔS_conf_ = S_conf_^Ub^ – S_conf_^D2^. (H) The segments with -TΔS more than (mean + SD) are colored purple on Fat10 domains. Higher values of -TΔS suggest increased conformational flexibility in these Fat10 regions compared to ubiquitin.

Tc’s were measured in ubiquitin and plotted against each residue in Figure 4B (fitting errors in Figure S11B). The loop between β1 and β2 had high negative Tc values in ubiquitin, which correlates with the lack of interactions and disorder propensity in this region^31^. Due to the low sequence similarity between Fat10 domains and ubiquitin, comparing the Tc values between individual residues of the two proteins is challenging. Instead, Tc values were averaged over the different protein segments and compared (Figure S11C). The difference in averaged Tc values between the N-terminal domain and ubiquitin indicates that Fat10 hbonds are weaker in the β1β2 region and the α2 loop (Figure 4C, 4D). Similarly, the β1β2 regions and α1 have weaker hbonds in the C-terminal domain. The Tc measurements of the isolated Fat10 domains yielded similar results (Figure S11D-J).

To estimate the conformational entropy in Fat10, the backbone order parameters (S^2^) were measured by standard NMR relaxation experiments. The spin-lattice relaxation R1, spin-spin relaxation R2, and heteronuclear NOEs (hetNOE) were measured in Fat10 and ubiquitin (Figure S12). S^2^ values were calculated using the Lipari-Szabo Model-Free Analysis method and averaged over the protein segments. The difference in averaged S^2^ between Fat10 and ubiquitin was converted to conformational entropy –TΔS ^32, 33^ and plotted in Figure 4F-H. The higher value of -TΔS suggests higher dynamics and flexibility. The β1β2 and α2 loops had high conformational entropy in the first Fat10 domain than ubiquitin (Figure 4F, 4G). In the second domain, β1β2, β3β4, and helix α1 were more entropic compared to ubiquitin, the most significant being α1 (Figure 4G, 4H). Overall, salt bridges increase interactions between α1 and β1β2 and between α1 and the α2 loop in ubiquitin. In their absence, these regions in Fat10 have weaker hbonds and higher conformational entropy.

### Fat10 increases substrate unfolding and degradation in cells

We then studied the degradation of Fat10 conjugated substrates in cells by a cycloheximide assay using GFP as the model. Fat10-conjugated GFP was degraded to 40% within 4 hours after cycloheximide treatment, much faster than ubiquitin-conjugated GFP (Figure 5A, S13A). It is unclear if the higher degradation rate is exclusively because the proteasome-targeting tag Fat10 degrades rapidly, thereby accelerating substrate degradation, or if it also affects the substrate’s thermodynamic stability and induces partially unstructured regions. To study changes in the substrate thermodynamic stability due to Fat10 in cellular conditions, a heterologous system has to be used where Fat10 substrates are not degraded. The bacterial Pup-proteasome system functions by a distinct mechanism^34^ that does not recognize or degrade the eukaryotic Ub/Fat10 conjugated proteins and can be used as a heterologous host. The CRABP1 protein was chosen as the model substrate. CRABP1 can be engineered to bind a fluorescent dye that is quenched in the native state but not in the unfolded state^35^ (Figure 5B). Cells expressing CRABP1 were treated with various urea concentrations. The difference in fluorescent signals between 0M and 3M Urea was lowest in apo CRABP1, higher in Ub-CRABP1, and highest in Fat10-CRABP1, suggesting the CRABP1 stability is lowest when conjugated to Fat10 (Figure S13B-D). ΔG_unfolding_ was 2.4 kcal/mol less in Fat10-CRABP1 than Ub-CRABP1 (Figure 5D), which suggested a significant increase in substrate unfolding by Fat10 conjugation.

**Figure 5.**
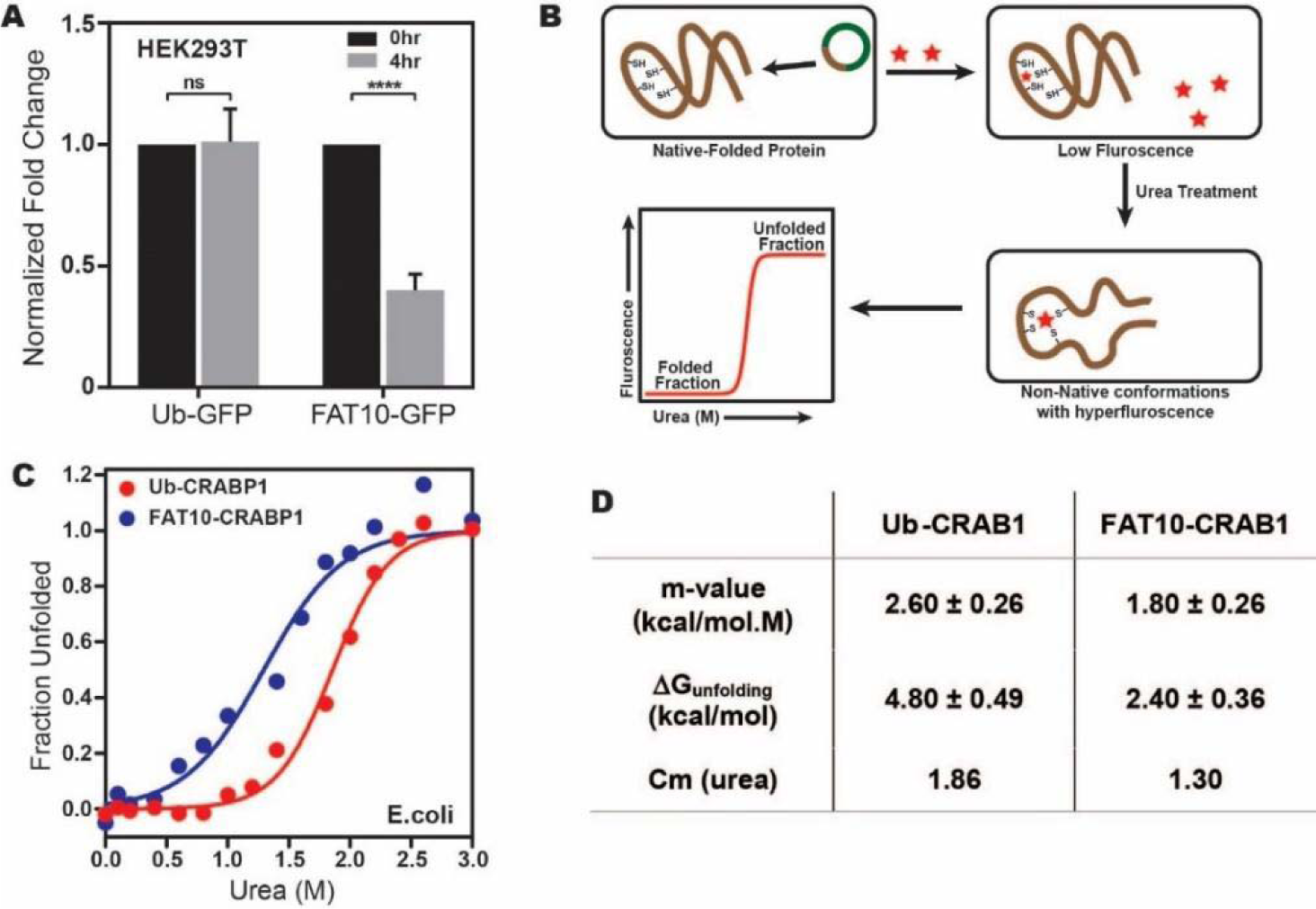
(A) Comparison of Ub-GFP and Fat10-GFP levels post 4hrs treatment with Cycloheximide (CHX) in HEK293T cells. (B) The schematic for studying the substrate stability in a heterologous cellular system, where a protein (CRABP1) capable of binding FIAsH-EDT2 dye was expressed in *E.coli* BL21 (DE3) cells as a substrate. The folded protein quenches the dye, while unfolded protein releases the quenching. (C) Fluorescence signals of Ub-CRABP1 and Fat10-CRABP1 were normalized to plot their denaturation curves. (D) Thermodynamic parameters of CRABP1 in cellular conditions when covalently bound to ubiquitin and Fat10, respectively.

### Fat10 induces partially unfolded forms in the substrate

In highly crowded cellular environments, proteins experience various physical interactions, such as the exclusion volume effect and nonspecific transient interactions. To study whether the substrate destabilization is solely due to Fat10 and not the above effects, the impact of Fat10 was studied in purified proteins. Producing purified Fat10 isopeptide conjugated substrates is technically challenging owing to the lack of a suitable Fat10 E3-substrate pair active under *in vitro* conditions. Hence, Fat10 was covalently conjugated at the N-terminus of a model ultra-stable substrate protein like the Cyan Fluorescent Protein (CFP) to mimic the N-terminal isopeptide conjugation ^36^ (Figure 6A). The free energy of unfolding CFP was calculated by measuring CFP fluorescence during chemical denaturation. CFP is a well-folded substrate with high thermodynamic stability and ΔG_unfolding_ = 11 kcal/mol. The change in unfolding energy of CFP is modest ΔΔG_unfolding_ = 0.2 kcal/mol when conjugated with ubiquitin (Figure 6B, S14A). The modest decrease in CFP stability correlates with the finding that ubiquitin conjugation is insufficient for the direct degradation of well-folded substrates and requires Cdc48^18^. Interestingly, ΔΔG_unfolding_ of CFP was significantly higher (3 kcal/mol) when covalently linked to Fat10, suggesting Fat10 destabilizes the substrate CFP 15-fold more than ubiquitin (Figure 6B, Figure S14A).

**Figure 6.**
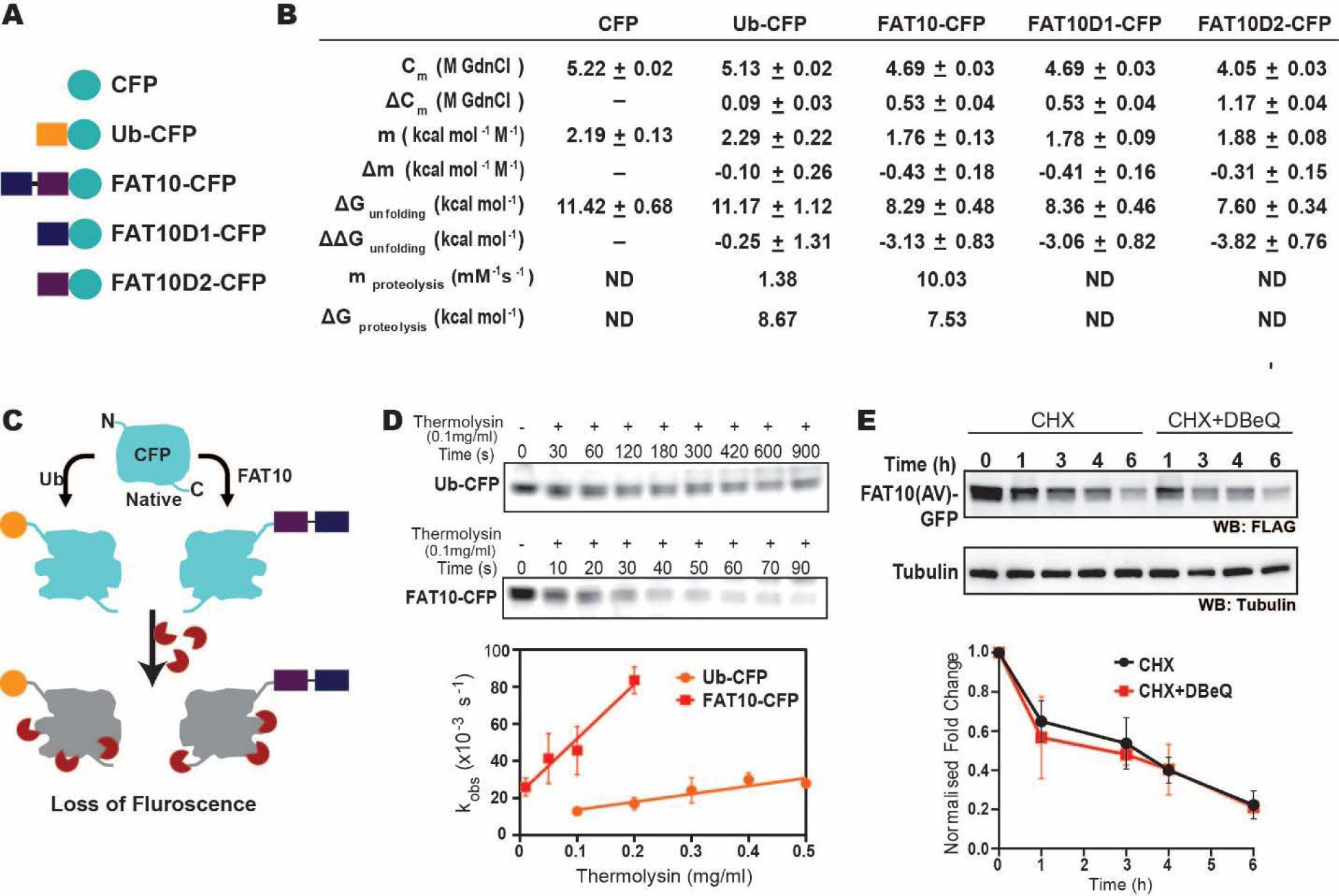
(A) This study used Different CFP versions, where CFP is fused at the C-terminal end of Ub, Fat10, Fat10D1, and FAT0D2. (B) A table with thermodynamic and proteolysis parameters of CFP bound to Fat10 domains is provided. (C) Native state proteolytic cleavage of Ub-CFP and Fat10-CFP using thermolysin. (D) Representative in-gel fluorescence image for native-state proteolysis of Ub-CFP and Fat10-CFP at 0.1 mg ml^-1^ thermolysin. k_obs_ of proteolysis are plotted for Ub-CFP and Fat10-CFP against various thermolysin concentrations. Error bars denote the standard deviation of the replicates (n=3). (E) *In-vivo* degradation of Fat10-GFP after cycloheximide treatment without/with p97 inhibitor (DBeQ). Tubulin is used as the loading control. The quantified level of FLAG-Fat10-GFP without/with inhibitor is given below.

Change in free energy could be due to the nonspecific or specific interactions between Fat10 and CFP ^37^. Specific interactions should depend on the complementary surface properties of Fat10 and CFP and should be affected if the Fat10 is truncated to smaller individual domains. CFP’s thermodynamic stability after N-terminal conjugation with the Fat10 domains (Figure 6A). The ΔΔG_unfolding_ was 3.0 kcal/mol and 3.8 kcal/mol when CFP is conjugated to Fat10D1 and Fat10D2, respectively (Figure 6B, S14B). The ΔΔG_unfolding_ due to the isolated domains is similar to the full-length Fat10, suggesting that instead of specific interactions, nonspecific interactions due to conjugation reduce CFP stability.

Fat10 may increase the partially unfolded regions in the substrate, which is critical to interact with the proteasome. The susceptibility of partially unfolded conformations to a typical protease thermolysin was measured in CFP, ubiquitin-CFP, and Fat10-CFP (Figure 6B-6D, Figure S14C-F). Although CFP was stable in the presence of protease, ubiquitin-CFP and Fat10-CFP were proteolyzed. Fat10-CFP proteolysis rate (k_obs_) was significantly more than ubiquitin-CFP at a given thermolysin concentration (Figure 6D). The change in observed proteolysis rates with different thermolysin concentrations (m_proteolysis_) was seven-fold higher in Fat10-CFP than in ubiquitin-CFP (Figure 6D). The values of m_proteolysis_ can be correlated to the free energy of proteolysis^38^. The free energy of proteolysis decreased from 8.7 kcal/mol for ubiquitin-CFP to 7.5 kcal/mol for Fat10-CFP (Figure 6B), indicating that Fat10 substantially increases the partially unstructured regions in CFP. Our results indicate that Fat10 has a more drastic effect than ubiquitin on substrate stability, suggesting that Fat10 substrates may be degraded independently of the unfoldases^15^. Degradation of Fat10-GFP was monitored in the HEK293T cells in the presence and absence of the Cdc48 inhibitor DBeQ (Figure 6E). Fat10-GFP degradation was unperturbed by DBeQ, suggesting that Fat10-modified substrates are degraded independently of Cdc48. Altogether, Fat10 significantly destabilizes substrates and creates partially unstructured regions for direct proteasomal degradation.

### Fat10 reduces local stability in the substrate

The mechanism of Fat10-induced disorder in the substrate is unclear. We used a combination of computational and experimental frameworks to detect changes in the substrate’s local stability. Ubiquitin was chosen as a well-folded small substrate as covalent interactions between Fat10 and ubiquitin have been reported previously^12^. MD simulations studied the impact of Fat10 on the kinetics of substrate ubiquitin unfolding at 450K. As a control, we used diubiquitin, where the first ubiquitin serves as the tag and the second ubiquitin is the substrate. The substrate ubiquitin in diubiquitin unfolded similarly to its free form (Figure S15A, C). However, the substrate ubiquitin conjugated to Fat10 domains unfolded faster (Figure S15B, D), suggesting that Fat10 domains increase the rate of substrate unfolding. Fat10 also enhanced fluctuations in the helix α1 and the loop near α2 in the substrate (Figure 7A). It increased the radius of gyration in substrate ubiquitin from 11 Å to 12 Å and created a broad RMSD profile starting from 2 Å to 3 Å, indicating a less compact substrate (Figure 7B). The D2 domain had a similar effect and created partially disordered states in the substrate (Figure 7A, 7B).

**Figure 7.**
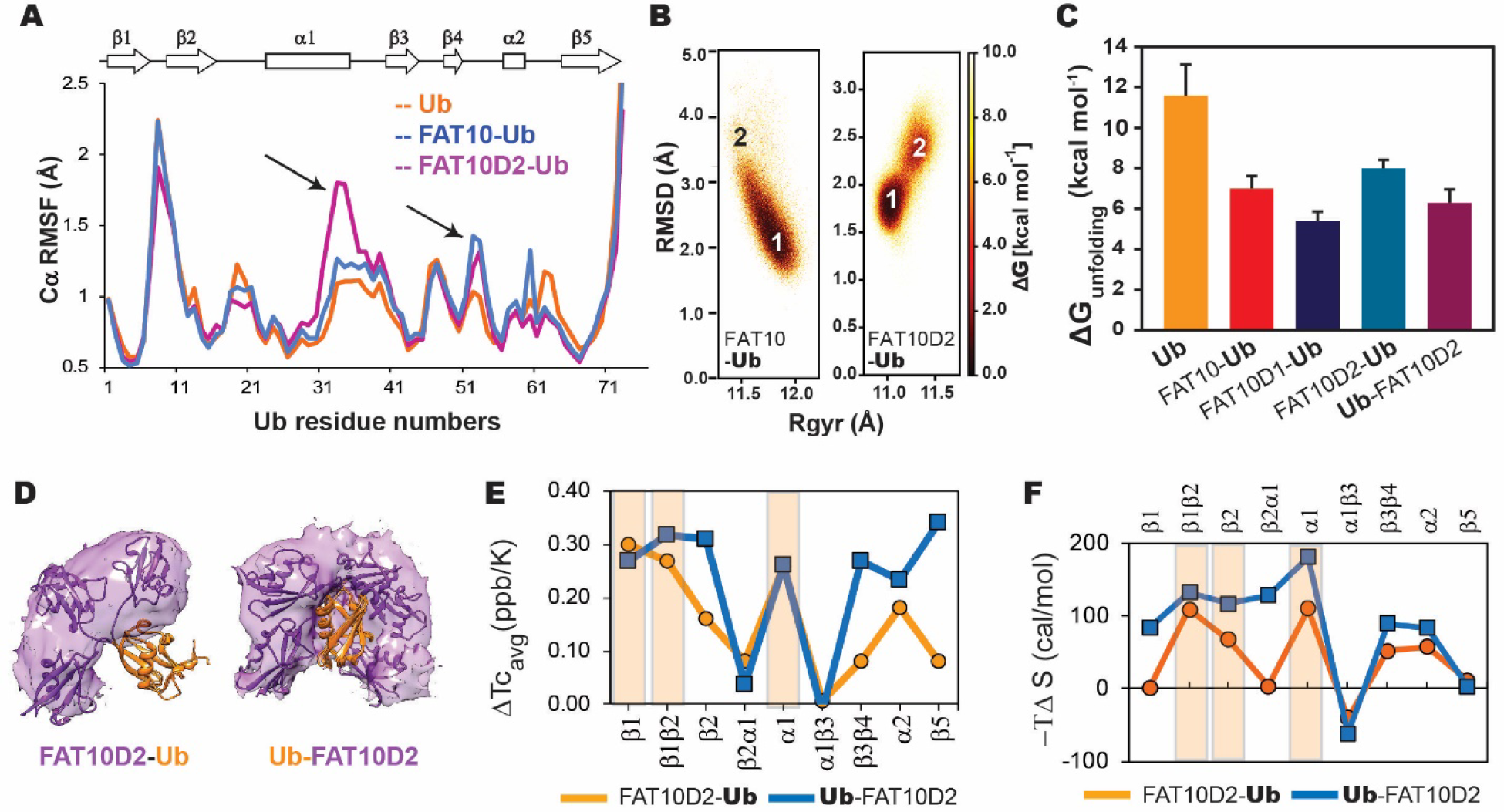
(A) Comparing Cα RMSF values of ubiquitin in monoubiquitin, Fat10-Ub, and Fat10D2-Ub. Black arrows denote regions with higher values in Fat10-conjugated ubiquitin than monoubiquitin. (B) The free energy landscape of Ub in Fat10-Ub and Fat10D2-Ub are plotted as a function of RMSD and radius of gyration (Rgyr) obtained from simulations across three replicas (3 x 2.5 μs) performed at 300K. (C) The free energy of unfolding ubiquitin in free form and covalently bound to Fat10 domains is plotted. (D) The occupancy of the D2 domain around Ub in the simulations is shown as a purple surface for Fat10D2-Ub and Ub-Fat10D2. A few structures in the simulation are superimposed and shown. (E) The difference in mean temperature coefficients between Fat10D2-Ub and free Ub is plotted for the various Ubiquitin segments. The orange line is ΔTc_avg_= Tc(Ub)_avg_ - Tc(Fat10D2-Ub)_avg_. The blue line is the same for Ub-Fat10D2, where ΔTc_avg_= Tc(Ub)_avg_ - Tc(Ub-Fat10D2)_avg_. A light orange box highlights the regions with high ΔTc_avg_ in Fat10D2-Ub. Ub-Fat10D2 has additional regions with ΔTc_avg_ values. (F) The difference in conformational entropy -TΔS of ubiquitin between free Ub and Fat10D2-Ub, where ΔS = S^Ub^ – S^Fat10D2-Ub^ was averaged for the various secondary structures and loops (orange). The same was plotted for Ub-Fat10D2 in blue. A light orange box highlights the regions with high -TΔS in Fat10D2-Ub.

Fat10’s effect on substrate stability was experimentally measured using Ub_45F/W_ mutant as a substrate, where the intrinsic tryptophan fluorescence reports the thermodynamics of substrate unfolding. Ub_45F/W_ is functionally active, and its thermodynamic stability is similar to ubiquitin ^39^. A single tryptophan in the N-terminal Fat10 domain is mutated to Phenylalanine (F) to not interfere with the fluorescence spectra (Figure S15E). Free energy of unfolding the substrate reduced from 11.7 kcal/mol to 7 kcal/mol when covalently bound to Fat10 (Figure 7C, Figure S15F-H). Individual domains of Fat10 had a similar effect on the substrate. C-terminal conjugation has a higher impact than at the N-termini (Figure 7C, Ub-Fat10D2 versus Fat10D2-Ub), suggesting that the conjugation site may regulate substrate stability.

To further understand substrate destabilization, we analyzed the tag-substrate and intra-substrate contacts. A significant number of tag-substrate contacts were formed between the D2 domain and the β1β2 and β4-α2-β5 regions in the substrate ubiquitin (Figure S16A, B). We then compared the long-range intra-substrate contacts in the free substrate and the Fat10-substrate conjugate (Figure S16C). The regions where intra-substrate contacts are disrupted overlapped with those where Fat10 contacts the substrate, suggesting that Fat10 interaction disrupts the intra-substrate contacts (Figure S16B, S16C). To study the higher destabilizing effect of Fat10 compared to ubiquitin, we noted that Fat10 has a higher exposed hydrophobic surface area (23%) than ubiquitin (13%) and can form more hydrophobic interactions with the substrate. Moreover, Fat10 domains are labile and sample partially unfolded forms, exposing their buried hydrophobic residues to enhance hydrophobic interactions with the substrate. We compared the intra-substrate long-range contacts between a linear diubiquitin molecule and Fat10-ubiquitin conjugate. The first ubiquitin serves as the tag in diubiquitin, and the second ubiquitin is the substrate. The number of intra-substrate contacts disrupted in the Fat10-substrate conjugate was higher than diubiquitin (Figure S17A, B), confirming the higher destabilization effect of Fat10.

Most intermolecular contacts were observed between the Fat10 D2 domain and the substrate, and few between the D1 domain and substrate. We conjugated the D2 domain to the N-terminal or C-terminal end of the substrate ubiquitin to study the significance of the conjugation site. The C-terminal tail in substrate ubiquitin is dynamic and explores a larger conformational space, providing greater flexibility for the tag to interact with the substrate (Figure 7D). Consequently, the intermolecular contacts between the tag and substrate were higher (Figure S16A, S17C, S17D). In addition, a more significant number of the intra-substrate contacts were disrupted (Figure S17A, E, F), which corroborates the reduced substrate stability upon C-terminal conjugation (Figure 7C, Ub-Fat10D2 versus Fat10D2-Ub). These results highlight that conformational dynamics at the conjugation site increase tag-substrate contacts and decrease substrate stability.

We used NMR spectroscopy to monitor the Fat10-induced changes in the substrate’s local stability. Since the destabilizing effect of Fat10 and the D2 domain are similar, and to avoid overlap in the NMR spectra, we studied the conjugates where the D2 domain is conjugated to substrate ubiquitin at its N- and C-termini (Fat10D2-Ub and Ub-Fat10D2). ^15^N-edited HSQC spectra of the conjugated proteins showed well-dispersed backbone amide resonances, confirming that they are folded (Figure S18). The Tc values were measured to measure Fat10’s effect on the strength of hbonds in the substrate (Figure S18C). Differences in averaged Tc values between Fat10D2-ubiquitin and free ubiquitin show that the hbonds in the strands β1β2, helix α1, and the α2 loop in the substrate ubiquitin are weakened (Figure 7E). Additional hbonds are destabilized in β3β4 and β5 when Fat10 is conjugated at the C-terminus, which is commensurate with its lower ΔG_unfolding_. Experimental NMR Tc measurements correlate excellently with the changes in local intra-substrate contacts observed in the MD simulations (Figure 7E and S17E-F). The substrate’s conformational entropy was quantified from order parameters (Figure S19). Conjugation at the N-terminal increases entropy at the β1β2 loop, β2, and α1 in the substrate (Figure 7F). Entropy increases further in these regions when D2 is conjugated at the C-terminus. In addition, the α2 loop and β3β4 regions become more entropic. Overall, the Fat10 domain contacts the substrate at multiple regions to weaken its local stability and increase disorder. The intrinsic flexibility of the conjugation site regulates Fat10’s impact on substrate stability.

The substrate may also influence the stability of the proteasome-targeting tag. The Fat10D2 tag is more compact (low Rog) in free form than in the tag-substrate conjugate (Figure S20A). Moreover, the Fat10D2 tag unfolds faster when conjugated with the substrate (Figure S20B), suggesting a thermodynamic coupling between the tag and the substrate. The tag-substrate contacts are greater when the tag is conjugated to ubiquitin C-termini (Figure 7D). Consequently, the tag unfolded faster when conjugated to the substrate C-termini (Figure S20C), suggesting that the tag stability also depends on the conformational flexibility of the conjugation site.

## Discussion

Posttranslational modification with proteasome-targeting tags is necessary but insufficient for protein degradation. The substrate/proteasome engagement and the substrate unfolding are rate-limiting steps for degradation^40, 41^. Global disorder, topology, local regions with a high disorder, and biased sequences in the substrate are critical factors that regulate proteasomal degradation^9^. Whether the structural flexibility of the proteasome-targeting tags impacts the substrate’s degradation rate is unclear. We report that the proteasome-targeting tag Fat10 has a malleable structure that samples multiple partially unfolded forms at physiological temperature. Lack of strong long-range electrostatic interactions between the central helix α1, the β1β2 strands, and the α2 loop, creates the flexible fold that provides weak resistance to mechanical unfolding. These properties of Fat10 expedite its unfolding, degradation, and degradation of Fat10 substrates, underlying the role of structural plasticity of proteasome-targeting tags in regulating protein degradation rate. Our data correlate well with the previous observations that substituting Fat10 domains with ubiquitin impedes proteasomal degradation^15^.

Fat10’s impact on the substrate structure and dynamics is noteworthy, as it destabilized various substrates in both *in-vitro* and cellular conditions. We chose two ultra-stable proteins, CFP and Ub, whose melting points are above 90°C. Fat10 reduced the unfolding energies of both substrates by 3-5 kcal/mol, which was considerably higher than ubiquitin. Our simulations suggested that Fat10 makes several nonspecific intermolecular contacts with the substrate to perturb the intra-substrate contacts. Competition between inter- and intra-molecular interactions has been recently highlighted, where critical salt-bridge interactions at protein-protein interfaces are “stolen” by new posttranslational modifications within the interacting proteins ^42, 43^. A similar mechanism may disrupt critical intra-substrate contacts by substrate-tag collision. Intriguingly, the substrate reciprocally increases partially unfolded regions in Fat10. Thermodynamic coupling between the substrate and tag functions in tandem to increase partially unfolded regions in the substrate-tag conjugate and accelerate its degradation.

Increased enthalpy and conformational entropy suggest local order-to-disorder transitions in proteins, giving rise to partially unfolded forms. We have measured the Tc values to estimate hbond strength, where higher ΔTc values indicate reduced hbond strength and increased enthalpy. Entropy was calculated from NMR relaxation order parameters. The increase in local enthalpy and entropy values identified Fat10-induced local disorder in the substrate, which agreed well with the computational data. An intriguing question is whether the chemical environment at the Fat10 conjugation site can regulate substrate stability. Our results show that conjugation at disordered regions allows greater conformational space for Fat10, increasing its collisions with the substrate and reducing substrate stability. Conjugation between two proteins can destabilize their folded state, stabilize their unfolded state, or both^37, 44^. Our NMR experiments have exclusively studied Fat10’s effect on the substrate’s folded state, and further studies are required to reveal its impact on the substrate’s unfolded state.

Posttranslational modification with ubiquitin destabilizes substrates ^19, 45–47^. However, the proteasomal degradation pathways of ubiquitin and Fat10 substrates are distinct. Post cleavage by proteasome deubiquitinases, ubiquitin’s impact on the substrate is unsustained. Hence, few substrates may not interact with the proteasome ATPases and escape degradation^9^. Fat10 remains conjugated to the substrates, and its effect on the substrate persists until translocation to the ATPases. Since Fat10 is degraded along with the substrate, its malleability directly influences the substrate degradation rate. While the N-termini of ubiquitin is rigid, a short N-terminal disordered region in Fat10 is cooperative to its degradation^15^. Moreover, Fat10 and ubiquitin’s impact on the substrate structure and energetics are distinct. Fat10 domains have a greater exposed hydrophobic surface than ubiquitin, which can induce more nonspecific substrate-tag collisions. Fat10’s ground state is in equilibrium with partially unfolded forms with exposed hydrophobic patches, further enhancing interactions with the substrate. Together, these structural and thermodynamic properties of Fat10 create substantially higher disorder in substrate than ubiquitin. These mechanistic differences in the pathways explain the unfoldase(Cdc48)-independent rapid degradation of Fat10 substrates. In hypoxic conditions, however, the 20S proteasome degrades ubiquitin conjugated to disordered substrates^48^. Since ubiquitin is uncleaved from the substrate, the mechanistic differences between the Fat10 and ubiquitin for 20S proteasomal degradation is an intriguing question for further investigation.

The proteasome degrades 80-90% of cellular proteins^49^, and understanding its activity has broad implications for our understanding of cellular response mechanisms to stress and infection. The immune cells undergo major reprogramming of signaling and antigen presentation during inflammation, and the proteasome plays an essential role in this process^50^. The fat10-proteasome pathway presents an efficient mechanism to upregulate protein degradation during inflammation. Intriguingly, it also presents an alternative for designing therapeutics by controlled protein degradation. Understanding the physical and thermodynamic effects of Fat10 modification is essential to create a model of how Fat10 serves as a rapid proteasomal degradation signal and engineering it for therapeutics. We measured the parameters of thermodynamic stability and degradation of Fat10 and Fat10 conjugated substrates. Our results suggest a loosely packed fold in Fat10 that is optimal for conjugation to substrates but unfolds effortlessly, allowing rapid protein degradation. It creates a significant disorder in the substrate for effective proteolysis. Fat10’s ductile fold is designed to function as an efficient proteasomal degradation signal that is activated as an inflammatory response.

## Acknowledgments

This work was supported by the Tata Institute of Fundamental Research, Department of Atomic Energy, Government of India, under project identification no RTI 4006. The NMR data were acquired at the NCBS-TIFR NMR Facility, supported by the Department of Atomic Energy, Government of India, under project no RTI 4006. The NMR facility is also partially supported by the Department of Biotechnology, India, under project number dbt/pr12422/med/31/287/2014. H.N. acknowledges a fellowship from UGC-CSIR, India. A.R. and P.D. acknowledge scholarships from the Department of Biotechnology, India.

## Authors Contributions

H. N., A.R., and R.D. designed the study. H.N. carried out experiments and data analysis. A.R. carried out simulations and data analysis. P.D. carried out ubiquitin denaturation experiments. S.R. carried out cell biology experiments. R.D. supervised the project. H.N. and R.D. wrote the initial manuscript draft, and all authors contributed to the final draft.

