## Supporting Information for "Plasticity of the proteasome-targeting signal Fat10 enhances substrate degradation"

Hitendra Negi[1,2], Aravind Ravichandran[1,2], Pritha Dasgupta[1], Shridivya Reddy[1#], and Ranabir Das[1*]

[1]National Center for Biological Sciences, TIFR, Bangalore, India

[2]SASTRA University, Thirumalaisamudram, Thanjavur, India

*Electronic address:

[#]Currently at Institute for Stem Cell Science and Regenerative Medicine (inStem), Bangalore, India

**Contents**

1. Text providing methods of Cloning, Protein Purification, Structure-based Phylogenetic analysis, Molecular Dynamics simulations, Far-UV Circular Dichroism (CD) measurements, NMR relaxation measurements
2. Supporting Figures S1-S20
3. Legends for the movies of Molecular Dynamics simulations S1 and S2
4. References for the text.

### Methods and Materials

#### Cloning

A *Fat10* gene (Life Technologies) was constructed where C7, C9, C160, and C162 were mutated to alanine. The gene was then sub-cloned in a pET3a vector tagged with 6xHis using NdeI and BamHI restriction sites and was used in the *in-vitro* experiments. The N-terminal domain (Fat10D1: 1-83aa with the GG motif at the C-terminal end)) and the C-terminal domain (Fat10D2: 84-165aa) were PCR amplified and further cloned into pET3a and pET14b vector, respectively. Fat10D1 was tagged with 6xHis, whereas Fat10D2 was tagged with 6xHis-scSMT. A precision protease site was introduced between the scSMT tag and Fat10D2.

The CFP plasmid was a generous gift from Dr. Aakash Gulyani, which was further sub-cloned into a pET3a vector using NdeI and BamHI restriction sites with a 6xHis tag at its N-terminus. CFP chimeric constructs Ub-CFP, Fat10-CFP, Fat10D1-CFP, and Fat10D2-CFP were cloned using overlapping PCR(1), where the amplified product of CFP was inserted at the C-terminus of the genes. For the ubiquitin chimeric constructs, Fat10D2 and Fat10D1 were swapped by ubiquitin to make Fat10D1-Ub and Ub-Fat10D2 constructs, respectively. The Fat10D2-Ub was created by first sub-cloning Fat10D2 in the pGEX6P1 vector with GST tag and then fusing the PCR product of ubiquitin at the C-termini of Fat10D2. Similarly, ubiquitin was inserted at the C-terminus of Fat10 to create the Fat10-Ub construct. For the tryptophan-based melt studies, phenylalanine (F) at the $45^{th}$ position of ubiquitin was mutated to tryptophan (W) in Ub, Fat10-Ub, Fat10D2-Ub, and Ub-Fat10D2. The tryptophan residue at the $17^{th}$ position in Fat10 was mutated to phenylalanine in Fat10-$Ub_{F45W}$ and Fat10D1-$Ub_{F45W}$ by site-directed mutagenesis. All the clones were verified by the Sanger sequencing method.

3xFLAG-wtFat10 (wild type 1-165) was constructed from Life Technologies for the cellular experiments. The gene was further sub-cloned into pcDNA3.1/hyg(+) vector (a kind gift from Dr. Apurva Sarin, NCBS) using HindIII and BamH1 sites. 3xFLAG-wtFat10D1 (aa:1-81) and 3xFLAG-wtFat10D2 (aa:82-165) were further made from 3xFLAG-wtFat10 in pcDNA3.1. After confirming the 3xFLAG-wtFat10D1 clone, a C-terminal tail sequence CYCIGG (same as the C-terminal tail of wtFat10) was inserted. The GG residues in 3xFLAG-wtFat10-GG, 3xFLAG-wtFat10D1-CYCIGG, and 3xFLAG-wtFat10D2-GG were also mutated to non-conjugal AV residues to create 3xFLAG-wtFat10-AV, 3xFLAG-wtFat10D1-CYCIAV, and 3xFLAG-wtFat10D2-AV in the pcDNA3.1 vector. Mutations in Fat10 were performed by site-directed mutagenesis. The Ubk0GV-GFP construct was obtained from Addgene (#11932) (2) and further sub-cloned into pcDNA3.1 to get 3xFLAG-Ub-k0GV. For the cycloheximide assay, GFP chimeric constructs were made by inserting the PCR product of GFP at the C-terminus end of 1xFLAG-UbGV and 3xFLAG-wtFat10AV by PCR. For the *in-cell* protein stability assay, a pseudo wild type CRABP1 gene (with R131Q stabilizing mutation and tetra-Cysteine motif by replacing G102C, D103C substitution followed by further insertion of 2 Cysteine residues between $^{105}$P and $^{106}$K) (3) was constructed from Life Technologies and sub-cloned in pET3a vector with 6xHis tag. The PCR amplified product of CRABP1 was inserted at the C-terminus end of ubiquitin, and Fat10 constructs to make Ub-CRABP1 and Fat10-CRABP1.

**Protein Purification**

All the clones were expressed in *E. coli* BL21 (DE3) cells. Post-induction with 1 mM IPTG, the bacterial culture transformed with Fat10, Fat10D1, Fat10D2, Ub-CFP, Fat10-CFP, Fat10D1-CFP, and Fat10D2-CFP were grown at 18ºC for 16 hrs, lysed, and resuspended in lysis buffer (Buffer A: 50 mM Tris buffer pH 7.5, 500 mM NaCl, 20 mM Imidazole and 1 mM DTT). The supernatant was loaded onto IMAC (GE) pre-packed column and eluted with imidazole gradient using Buffer A and Buffer B (Buffer A with 500 mM Imidazole). The purified fractions were applied on a size exclusion chromatography on a Superdex 16/600 75pg column (GE Healthcare) pre-equilibrated with SEC buffer (Buffer C: 50 mM Tris buffer pH 7.5, 250 mM NaCl, with or without 1 mM DTT). All the ubiquitin proteins were also purified, as given above. GST-ppx-Fat10D2-Ub was purified using GST pre-packed column and eluted with 10-20 mM reduced Glutathione. The purified proteins His-scSMT-Fat10D2, His-scSMT-ppx-Fat10D2-CFP, and GST-ppx-Fat10D2-Ub, were further treated with GST-tagged Precision protease enzyme followed by reverse IMAC. The digested Fat10D2, Fat10D2-CFP, and Fat10D2-Ub were purified by size exclusion chromatography. Ubiquitin and SUMO1 proteins were purified using protocols published elsewhere (4, 5). For the NMR experiments, Fat10D1, Fat10D2, Fat10-Ub, Fat10D1-Ub, Ub-Fat10D2, and Fat10D2-Ub were grown in M9 media containing isotopic Ammonium chloride ($^{15}NH_4Cl$) and/or isotopic Glucose ($^{13}$C-Glucose). The purification protocol was the same as given above.

**Structure-based Phylogenetic analysis**

The structures of 10 available UBL PDBs were retrieved from RCSB PDB as tabulated in Fig. S1A. Crystal structures were cleaned by removing co-crystal structures, multiple domains, ions, and water. For NMR structures, the best representative model was used. For Fat10 and ISG15, individual UBL domains were isolated. RMSD matrix and structure-based sequence alignment were obtained for all structures based on multiple superimpositions using the MUSTANG tool (6). Further, the phylogenetic tree was plotted using aligned sequences in MEGA X (7).

**Molecular Dynamics simulations**

Structures for individual domains of Fat10D1 & Fat10D2 were retrieved from PDB ids 6GF1 and 6GF2, respectively. The full-length Fat10 structure was modeled using Swiss-Modeller with 6GF1 and 6GF2 as templates(8). Chain B of 6GF1 was further processed for simulation and called Fat10D1. All water and sulfate molecules were removed from the Fat10D1 crystal structure. 6GF2 was an NMR ensemble structure, and the best representative structure, Fat10D2, was further considered for simulation. For chimeric models, the individual domains of Fat10D1, Fat10D2 & ubiquitin were fused between the N-terminal of one domain and the C-terminal of the other using UCSF Chimera (9). These models were further simulated for 500 ns before using the final structure for unfolding simulations. The salt bridge mutants were designed using the Rotamer function in UCSF Chimera. Systems for Fat10D1, Fat10D2, Fat10, and ubiquitin were prepared with the LEaP program of Ambertools18. Systems were prepared in a cubic box of TIP3p water, with a minimum distance of at least 12 Å between solute atoms and the box edge. Counter ions were added to neutralize the system.

Parameters describing system topology were based on the Amber ff99SBildn force field (10). The systems were first relaxed by energy minimization using the Sander module of Amber18 in two stages. In the first stage, water molecules were minimized with restraint on protein, and then the entire system was minimized. The respective systems were then heated incrementally in NVT from 0K to 300K for 5 ns with positional restraints (20 kcal/mol/$Å^2$) on protein atoms. Further, system density was equilibrated for 5 ns in the NPT ensemble with positional restraints (20 kcal/mol/$Å^2$) on protein atoms. Further, four subsequent equilibration stages reduced the restraints on the backbone atom from 20 to 0 through a series of molecular dynamics simulations in an NPT ensemble for 400 ps each. The final production run was performed for 2.5 µs in NPT with three replicas. The distance cutoff for short-range nonbonded interactions was set to 1 nm. The particle mesh Ewald (PME) method was used to treat long-range electrostatic interactions. The SHAKE algorithm (11) was applied to constrain all bonds involving hydrogen atoms. Temperature was set to 300K using a Langevin thermostat, and pressure was maintained at 1 bar using the Berendsen barostat. Using the hydrogen mass repartitioning (HMR) scheme (12), the integration time step was set to 4 fs. Dynamics were propagated using the leapfrog integrator. Snapshots were saved every 40 ps giving 65200 conformations from a single run. A total of 7.5 µs (3*2.5 µs) data was pooled for further analysis.

The trajectory analyses were performed using the AMBER suite's CPPTRAJ module (13). The averages and standard errors were calculated using in-house scripts and were plotted using the R program. Native contacts were calculated using the native-contacts method in CPPTRAJ with a 7 Å distance cutoff. For backbone native contacts across secondary structure pairs, the native contacts were calculated, defining a 3.5 Å distance cutoff on backbone atoms. The data was averaged across ten individual simulations at 450 K to calculate the mean and standard error. The Free Energy plots obtained from bin populations of the 2-dimensional histograms obtained from binning Root Mean Squared Deviation (RMSD) and Radius of Gyration values using the formula:

$$\Delta G_i = -k\beta * T * \ln(N_i/N_0) \qquad (5)$$

where $\Delta G_i$ is the free energy value of bin i, kβ is Boltzmann's constant in kcal/mol*K, T is the temperature in K, $N_i$ is the population of bin i, and $N_0$ is the population of the most populated bin.

RMSD and Rgyr were calculated using rms and rog methods available in CPPTRAJ with their respective crystal structure as a reference. The data were obtained from three individual 2.5 μs repeats at room temperature. Rgyr represents the Radius of Gyration for the protein without loops. The loops were omitted during the Free Energy calculation to avoid false positives. Rog represents the Radius of Gyration for the entire protein. Principle Component Analysis was performed on Ub, Fat10D1, Fat10D2, and their respective salt bridge mutants. PCA was also performed for Fat10D1-Ub, Fat10D2-Ub, and Ub-Fat10D2 using the Bio3D package(14) in R(15). PDB format trajectory was further produced that interpolates between the most dissimilar structures in the distribution along PC1. To obtain contact maps, per-residue native and non-native contacts were calculated from trajectories, normalized, and plotted using GNUplot.

**Far-UV Circular Dichroism (CD) measurements**

The Far-UV CD data was collected using Jasco J-1500 spectropolarimeter in a 1mm cylindrical quartz cuvette, using a 1nm/sec scan speed from 200 -250nm with a digital integration time of 4s. Five scans at 25°C were averaged and plotted using the SIGMA plot after converting the mdeg to MRE (Mean Residual Ellipticity). Appropriate buffer scans were subtracted from the protein's scan.

**NMR relaxation measurements**

For the studies of the Fat10 dynamics, uniformly labeled $^{15}$N-Fat10, $^{15}$N-Fat10D1, $^{15}$N-Fat10D1-Ub, $^{15}$N-Fat10D2-Ub, $^{15}$N-Ub-Fat10D2, and $^{15}$N-ubiquitin were dialyzed in 50 mM Sodium phosphate buffer, 250 mM NaCl with pH 7.5. The NMR relaxation data were collected at 298K on an 800 MHz and 600 MHz NMR spectrometer (Bruker). Longitudinal (T1), Transverse (T2) time constraints, and hetero-Nuclear Overhauser Enhancement (hetNOE) experiments were carried out using standard pulse sequences in Bruker. For T1 measurements, data were recorded at the following relaxation delays: 0.004, 0.03, 0.06, 0.1, 0.15, 0.2, 0.4, and 0.8 sec. For T2 measurements, the relaxation delays were set to 0.004, 0.017, 0.035, 0.051, 0.068, 0.086, 0.102, 0.119, and 0.136 sec. A 5 sec pre-saturation was used in the $^{15}$N-Heteronuclear NOE experiment. The reference experiment was carried out with 10 sec delay without pre-saturation. The T1-relaxation delays for $^{15}$N-Fat10 were: 0.01, 0.025, 0.05, 0.075, 0.1, 0.2, 0.3 sec whereas, the T2 delay time was: 0.01, 0.02, 0.4, 0.06, 0.08, 0.1 sec and the hetNOE was recorded using standard pulse sequence in Bruker with 5 sec as mixing time. The hetNOE was calculated as the ratio of intensities $I_{sat}/I_{ref}$. The hetNOE error was calculated as (NOE err/NOE) = $[(I_{ref}\ err/I_{ref})^2+(I_{sat}\ err/I_{sat})^2]^{1/2}$, where the error of each intensity measurement is the rmsd noise of each plane. Order parameter ($S^2$) was calculated using T1, T2, and hetNOE experiments in RELAX software(16, 17) using the Lipari Szabo model-free approach where the $S^2$, correlation time ($\tau_e$), and $R_{ex}$ were obtained. The $S^2$ values were converted to backbone conformational entropy using the entropy meter approach(18, 19).

The temperature dependence of chemical shifts was measured for uniformly labeled $^{15}$N-Ub, $^{15}$N-Fat10D1, and $^{15}$N-Fat10D2 in 50 mM PO4 buffer with 250 mM NaCl at pH 7.5 on the 600MHz spectrometer. The temperature varied from 283K to 313K. The same was measured for $^{15}$N-Fat10, $^{15}$N-Fat10-Ub, and $^{15}$N-ubiquitin at the 800MHz spectrometer. Similar experiments were carried out in 50 mM PO4 buffer with 250 mM NaCl at pH 6.5 for the proteins $^{15}$N-Fat10D1-Ub, $^{15}$N-Fat10D2-Ub, and $^{15}$N-Ub-Fat10D2. The temperature varied from 283K to 323K. The chemical shift of water was used for reference, calibrated using temperature-independent 4,4-dimethyl-4-silapentane-1-sulfonic acid (DSS) signal. All the NMR HSQC experiments were processed by NMRpipe and analyzed in SPARKY. The chemical shifts in the $^1$H-dimension were analyzed using linear regression by MATLAB. The linear fit between the $\Delta\delta^{NH}$ and temperature provided the temperature coefficient of a residue, Tc = ($\Delta\delta^{NH}/\Delta T$). The Residual Sum Square (RSS) was calculated as RSS = $\sum_i^N (y_i - f(x_i))^2$, where $y_i$ is the i$^{th}$ measured value and $f(x_i)$ is the fitted value of $y_i$.

**A**

**List of Ubiquitin-Like Modifiers (ULMs)**

| ULMs | Uniprot ID | PDB ID |
|---|---|---|
| Ub | - | 1UBQ |
| NEDD8 | Q15843 | 1NDD |
| SUMO1 | P63165 | 1A5R |
| SUMO2 | P61956 | 1WM2 |
| SUMO3 | P55854 | 1U4A |
| UFM1 | P61960 | 1WXS |
| FAT10D1 | O15205 | 6GF1 |
| FAT10D2 | O15205 | 6GF2 |
| ATG12 | O94817 | 4NAW |
| ISG15 | P05161 | 1Z2M |

**B**

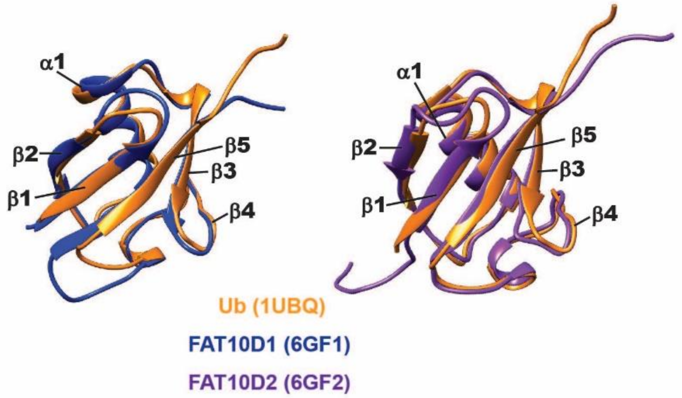


**C**

**RMSD (Å) between different ULMs**

| | SUMO1 | SUMO2 | SUMO3 | Ubiquitin | FAT10D1 | FAT10D2 | ISG15D1 | ISG15D2 | NEDD8 | UFM1 |
|---|---|---|---|---|---|---|---|---|---|---|
| SUMO1 | --- | | | | | | | | | |
| SUMO2 | 2.4 | --- | | | | | | | | |
| SUMO3 | 2.4 | 2 | --- | | | | | | | |
| Ubiquitin | 2.7 | 1.4 | 2 | --- | | | | | | |
| FAT10D2 | 2.8 | 1.8 | 2.1 | 1 | --- | | | | | |
| FAT10D1 | 2.8 | 1.6 | 1.9 | 1 | 1.1 | --- | | | | |
| ISG15D1 | 2.9 | 2.2 | 2.6 | 1.8 | 1.8 | 1.8 | --- | | | |
| ISG15D2 | 2.8 | 1.4 | 2 | 0.9 | 1.4 | 1.2 | 2 | --- | | |
| NEDD8 | 2.8 | 1.7 | 2.1 | 0.6 | 1 | 1.1 | 1.7 | 1.2 | --- | |
| UFM1 | 3.5 | 2.7 | 3 | 2.9 | 3.1 | 3 | 3.2 | 2.6 | 2.8 | --- |
| ATG12 | 3.2 | 2.2 | 2.5 | 2.2 | 2.3 | 2.3 | 2.6 | 2.2 | 2.3 | 3.1 |

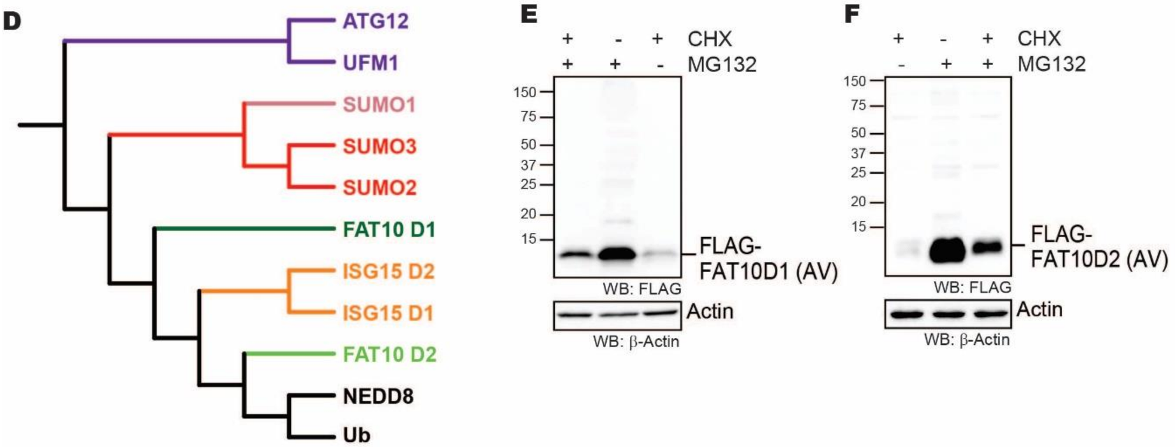


Figure S1. (A) List of Ubiquitin-Like Modifiers (ULMs) with their Uniprot and PDB ids used in making the phylogenetic tree. (B) The structure of Fat10D1 (6GF1) (Blue) and Fat10D2 (6GF2) (purple) is superimposed onto the structure of ubiquitin (1UBQ) (orange). (C) Table represents RMSD (Å) between different ULM pairs used to make phylogenetic trees. (D) Phylogenetic tree of 11 ULM based on their RMSD across the available structure in PDB. (E) HEK293T cells were transfected with FLAG-Fat10D1(AV) and treated with/without Cycloheximide and proteasomal inhibitor MG132 for 6 hours. The lysates were separated on SDS page gels and blotted with anti-FLAG antibodies. (F) The cells were transfected with FLAG-Fat10D2(AV), then treated and probed as in (D).

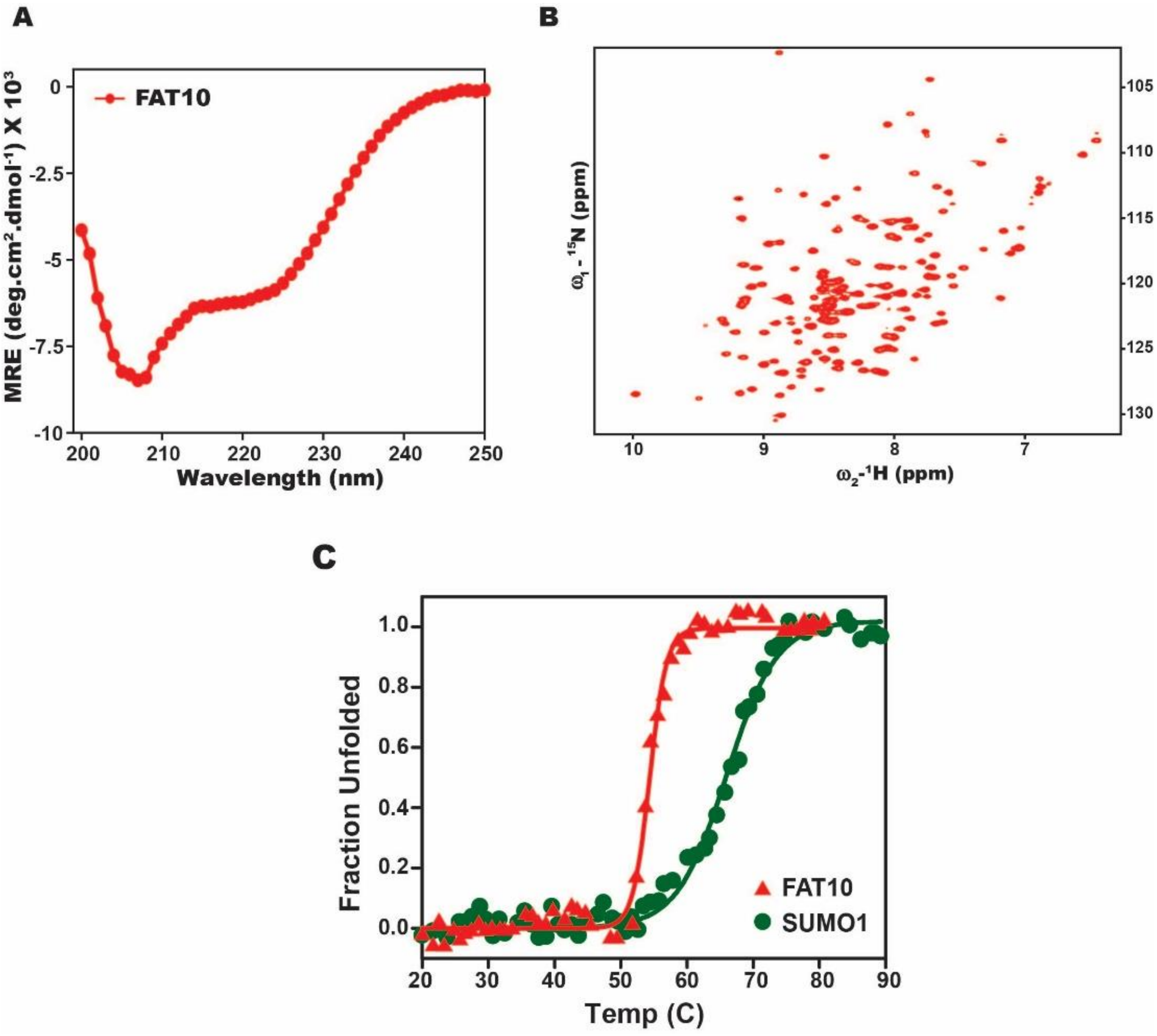


Figure S2. (A) Far-UV CD spectra and (B) $^{15}N$-$^{1}H$ HSQC spectra of purified Fat10. (C) The thermal melt curve of Fat10 and SUMO1, where the change in ellipticity is normalized and plotted against temperature.

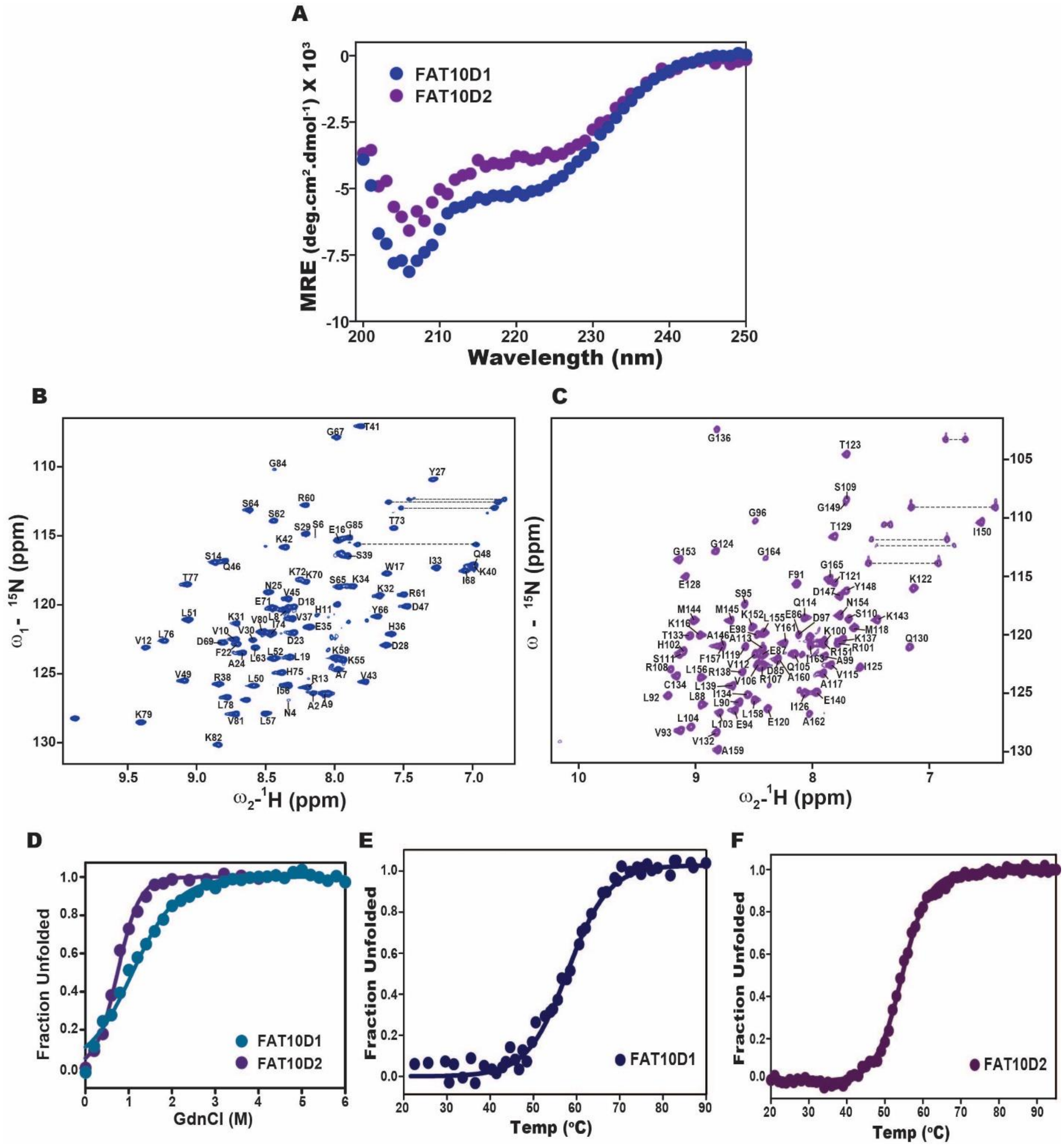


Figure S3. (A) Far-UV CD spectra of purified Fat10D1 (Blue) and Fat10D2 (Purple). (B) 2D $^{15}N$-$^{1}H$ HSQC spectra with peak assignment of Fat10D1and (C) Fat10D2. (D) The GdnCl melt curves of Fat10D1 and Fat10D2. The thermal melt curve of (D) Fat10D1 and (E) Fat10D2, where the change in ellipticity is normalized and plotted against temperature.

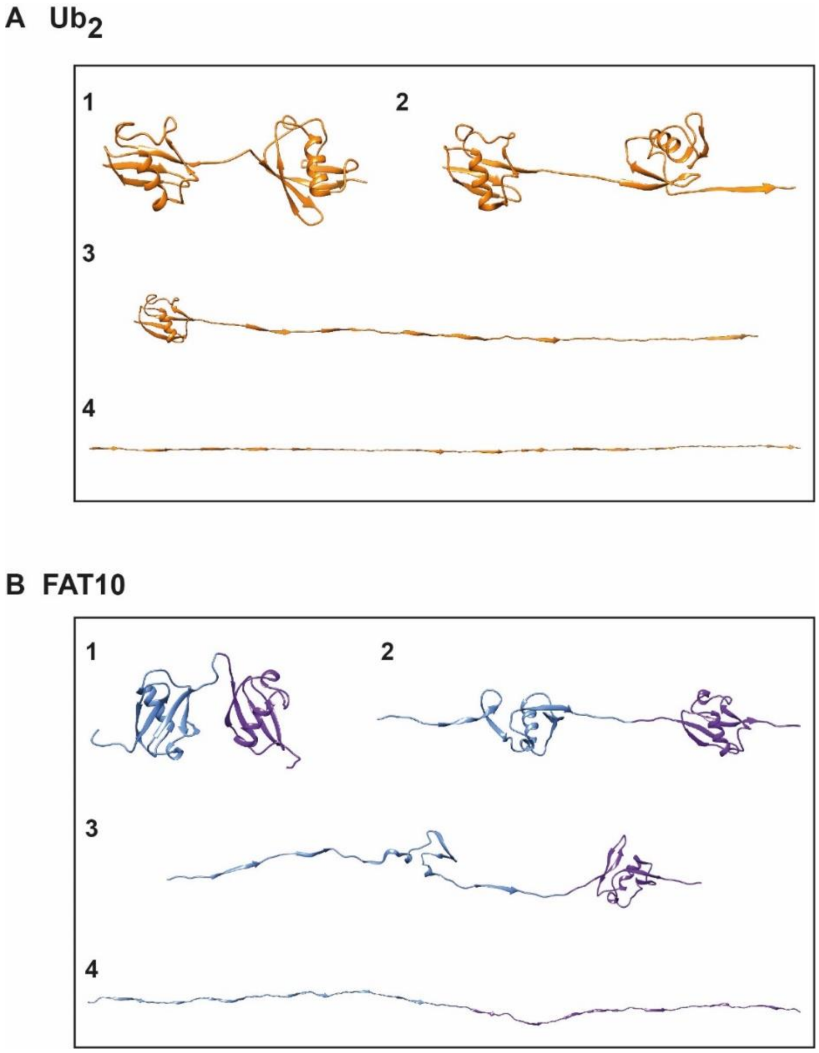


Figure S4. The intermediates observed during Adaptive Steered molecular dynamics of (A) $Ub_2$ and (B) Fat10.

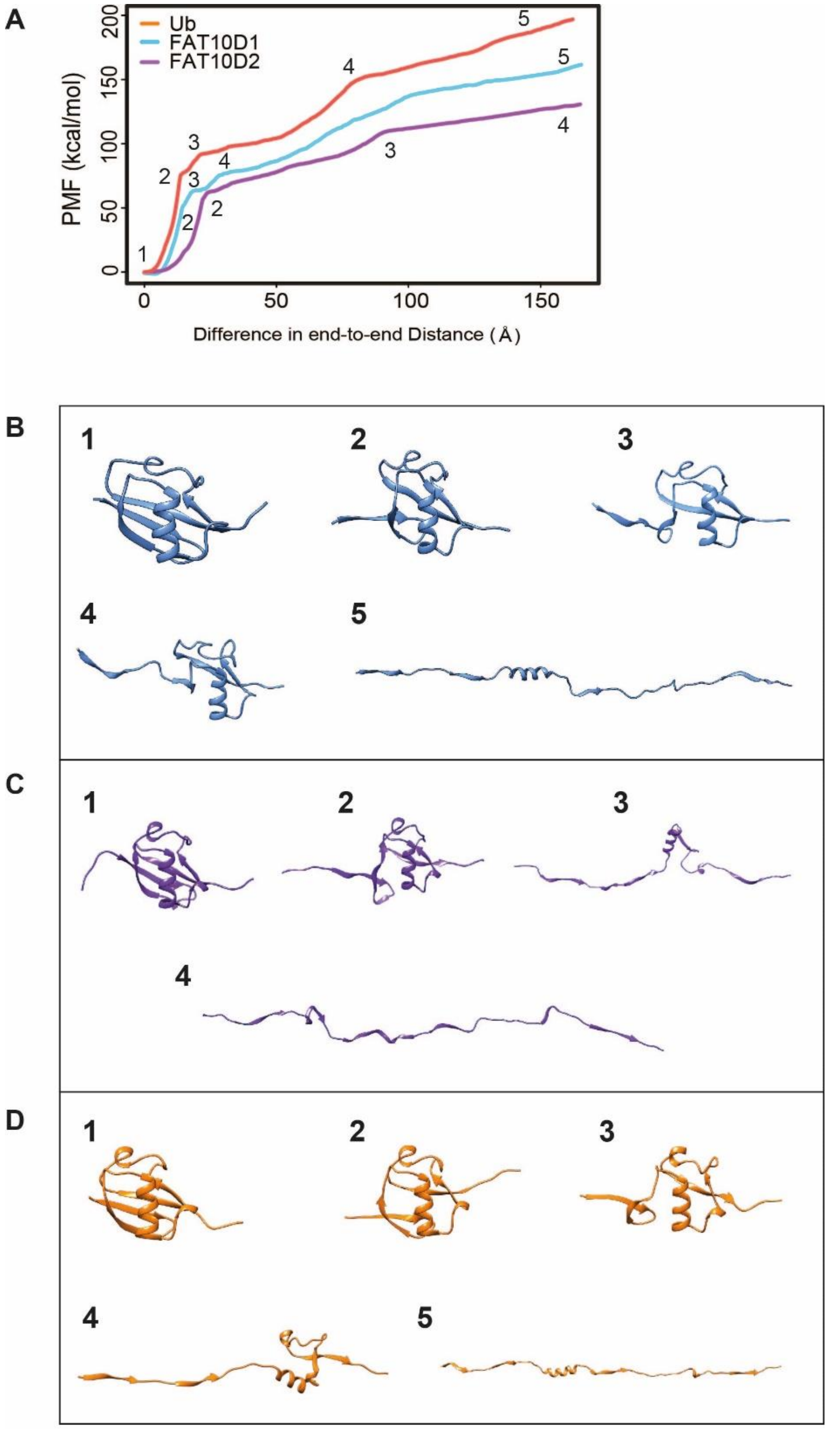


Figure S5. (A) Adaptive Steered molecular dynamics of Ub, Fat10D1, and Fat10D2. The potential Mean Force (PMF) is plotted against the normalized end-to-end distance. The pulling velocity is 1 Å/ns. Each unfolding event is marked by a number whose corresponding conformation is given in (B) Fat10D1, (C) Fat10D2, and (D) Ub.

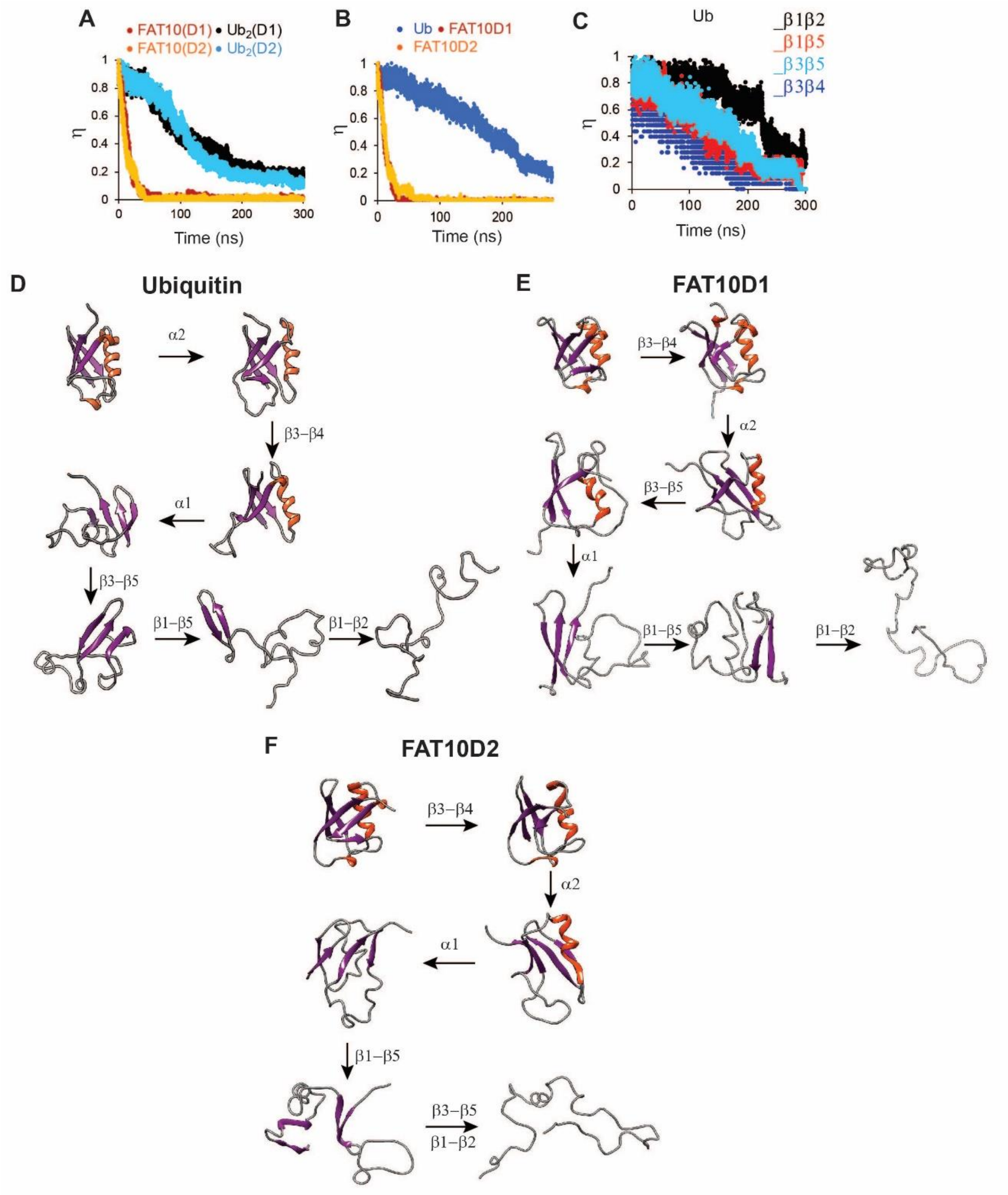


Figure S6. Molecular dynamics simulations of Fat10, di-ubiquitin ($Ub_2$), Ub, and individual domains D1/D2 in Fat10 were performed at 450K. (A) The fraction of native beta-sheet backbone hydrogen bonds is defined as ($\eta$) and plotted for Fat10 and $Ub_2$ against time. (B) is the same as (A) measured for individual domains Fat10D1, Fat10D2, and Ub. (C) The fraction of native pairwise backbone hydrogen bond is defined as ($\eta$), which is plotted for β1 to β5 of Ub. The intermediate structures of the unfolding pathway in (D) Ub, (E) Fat10D1, and (F) Fat10D2 were inferred by analyzing the fraction of backbone hbonds from ten replicas in the simulations.

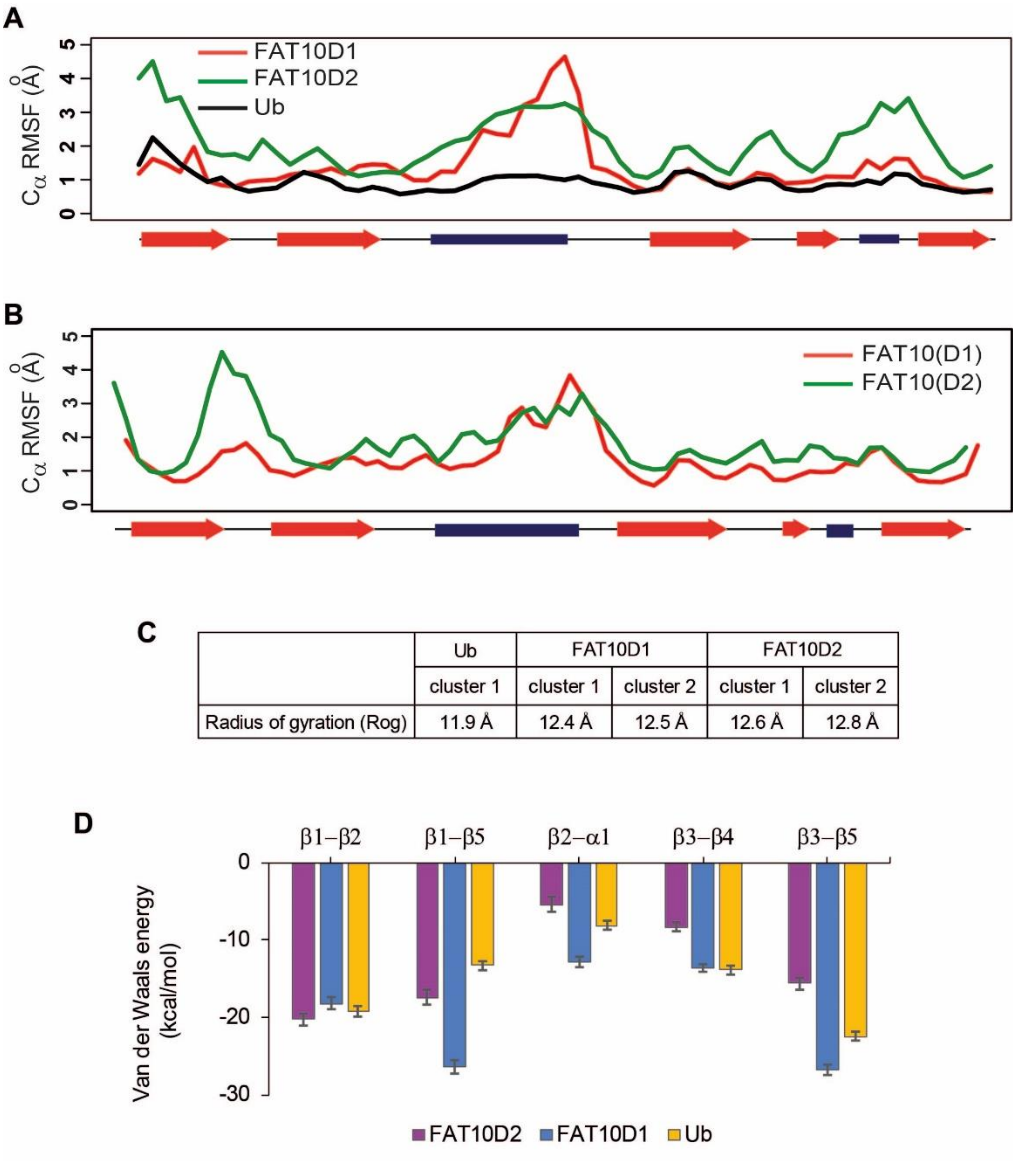


Figure S7. (A)The Cα root mean square fluctuations (RMSF) are plotted against residue numbers for the isolated domains Fat10D1, Fat10D2, and ubiquitin in the 300K simulations. (B) The Cα RMSF is plotted for the two domains in full-length Fat10. (C) The radius of gyration of the entire protein (Rog) for the minimas in Ub, Fat10D1, and Fat10D2 are provided. The Rgyr values do not account for long loops in the protein, while the Rog values include the complete protein. (D) The mean Van der Waals energy of interactions between different pairs of secondary structures obtained from 300K simulations is plotted for Ub, Fat10D1, and Fat10D2.

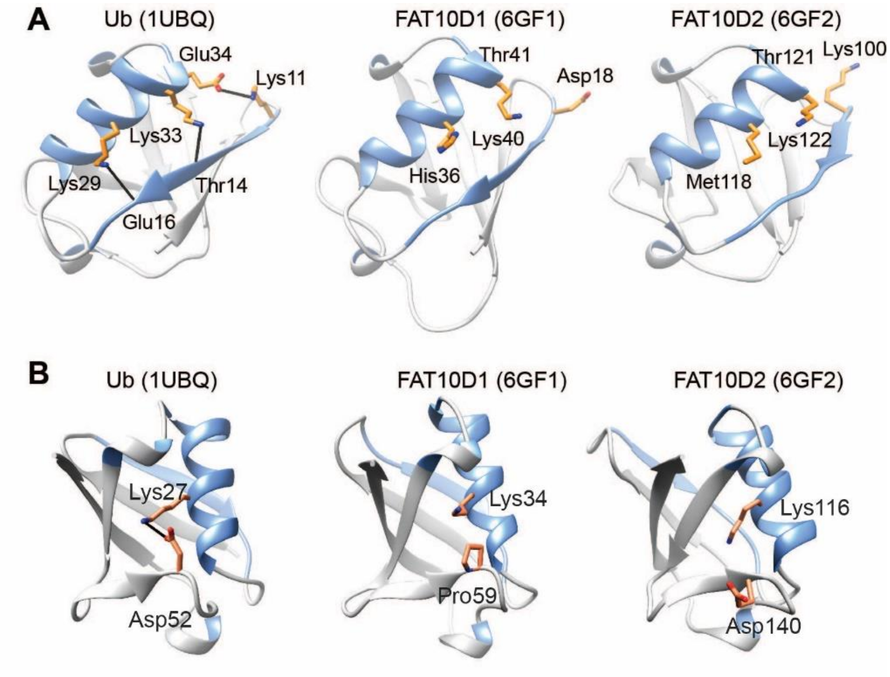


Figure S8. (A) The β2-α1 electrostatic interaction are mapped on Ub, Fat10D1, and Fat10D2 structures. The β2 and α1 are colored blue. The hydrogen bonds and salt bridges are shown as black lines. A Lys11-Glu34 salt bridge and two hydrogen bonds (Lys33-Thr14 and Lys29-Glue16) are observed in ubiquitin but absent in Fat10D1 and Fat10D2. (B) A K27-D52 salt bridge is observed in ubiquitin between the α1 and the α2 loop. A black line shows the salt bridge. In Fat10D1, K34 is present in α1 at the position analogous to K27. However, a proline residue P59 is present at a similar position to D52 in Ub. Consequently, the salt bridge is absent in Fat10D1. In the Fat10D2 structure, the D140 sidechain is facing away from the K116; hence, the salt bridge has not formed.

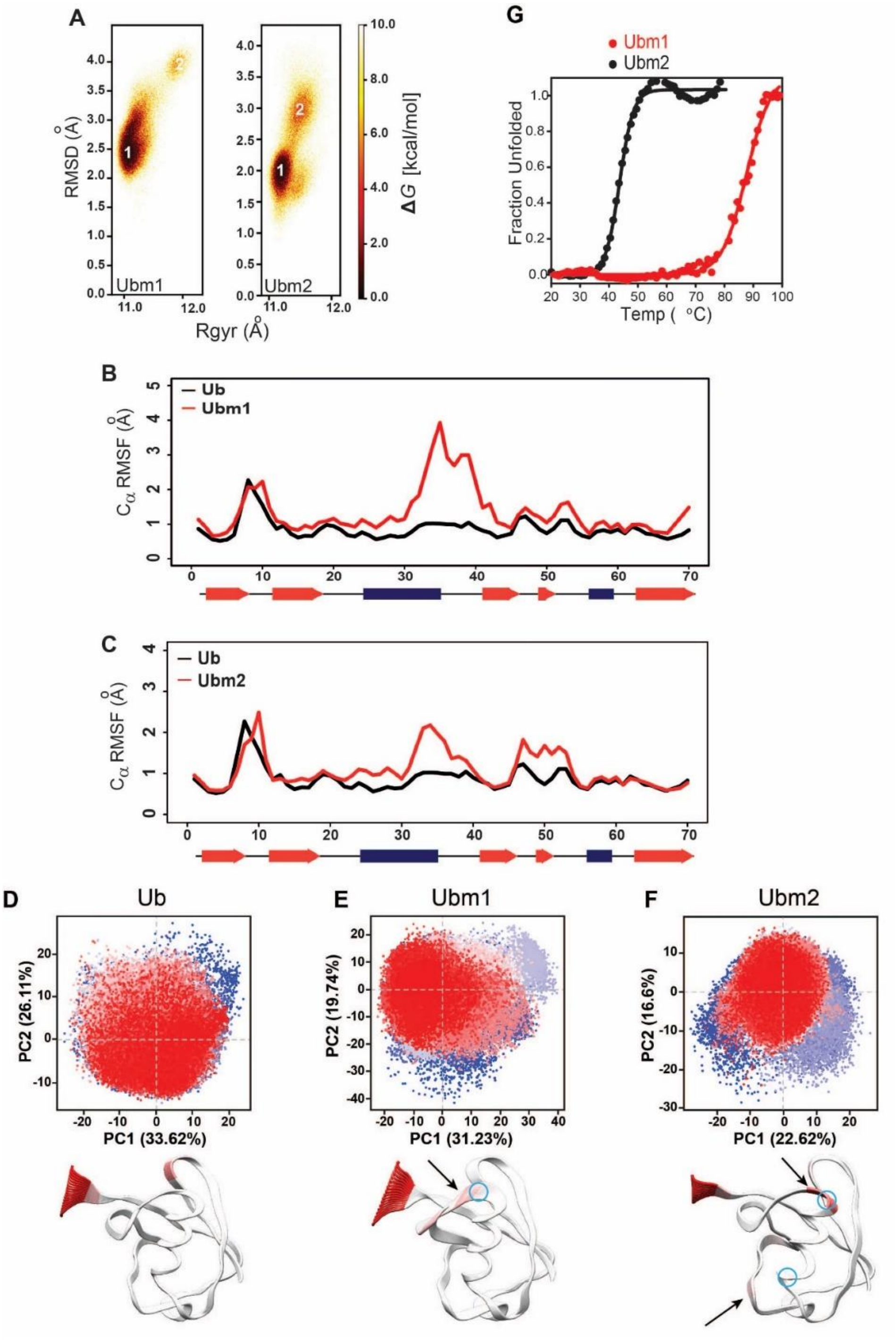


Fig S9. (A) The free energy landscapes of Ubm1 (E34K-Ub) and Ubm2 (E34K, K27D-Ub) are plotted as a function of RMSD and radius of gyration (Rgyr) obtained from simulations across three replicas (3 x 2.5 μs) performed at 300K. The minima from each cluster are numbered. (B) The Cα root mean square fluctuations (RMSF) are plotted against residue numbers for Ub and Ubm1. (C) The Cα RMSF values are plotted for Ub and Ubm2. (D) Distributions of different conformations sampled during the simulation were plotted onto principal component (PC) space for Ub and its mutants. The conformations are colored from blue to red in order of simulation time. The trajectory frames of ubiquitin on the PC axes for the first two components, PC1 and PC2. The variation observed along PC1 is mapped onto the structure of Ub. The thickness of the ribbon and color (red) is scaled by PC1 values. (E) and (F) are plotted similarly for the ubiquitin mutants Ubm1 and Ubm2, respectively. The black arrows indicate a gain in motions (fluctuations) with respect to wild-type Ub. The blue open circles indicate the site of substitution. (G) The thermal melt curve of Ubm1 and Ubm2, where the change in ellipticity is normalized and plotted against temperature.

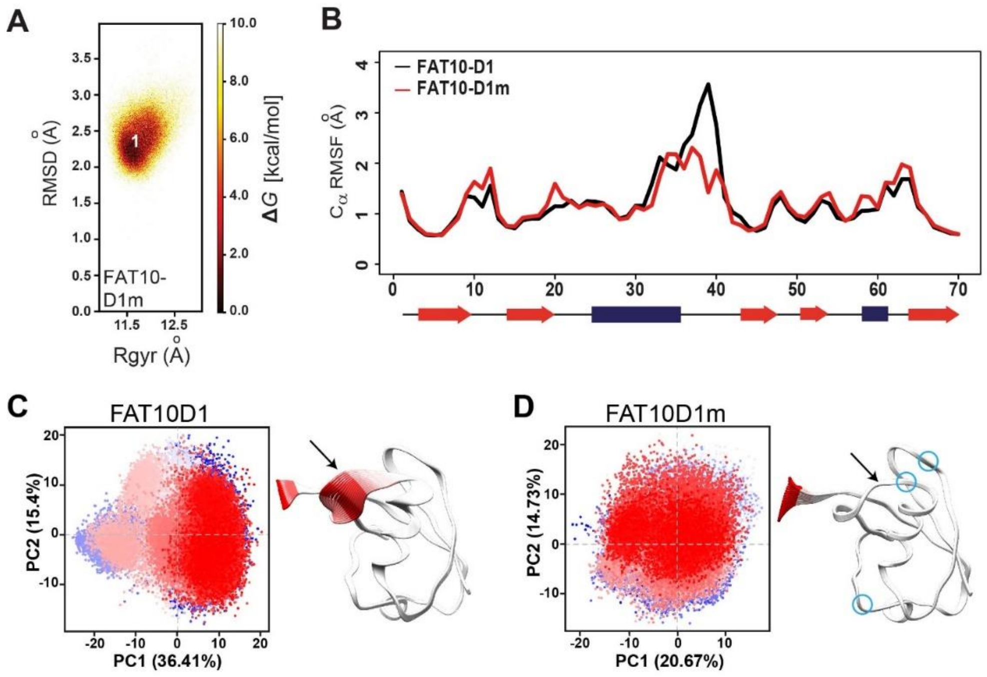


Fig S10. (A) The free energy landscape of Fat10D1m (D18K, T41E, P59D) is plotted as a function of RMSD and radius of gyration (Rgyr) obtained from simulations across three replicas (3 x 2.5 μs) performed at 300K. (B) The Cα RMSF values are plotted for Fat10D1 and Fat10D1m. (C)-(D) Distributions of different conformations sampled during the simulation were plotted onto principal component (PC) space for Fat10D1 and Fat10D1m. The conformations are colored from blue to red in order of simulation time. (C) Molecular dynamic frames of Fat10D1 on the PC axes for the first two components, PC1 and PC2. The variation observed along PC1 is mapped onto the structure of Fat10D1. The thickness of the ribbon and color (red) is scaled by PC1 values. (D) Similarly plotted for Fat10D1m. The arrows indicate loss or gain in dynamics (fluctuations) with respect to the wild-type protein. The blue open circles indicate the site of substitution.

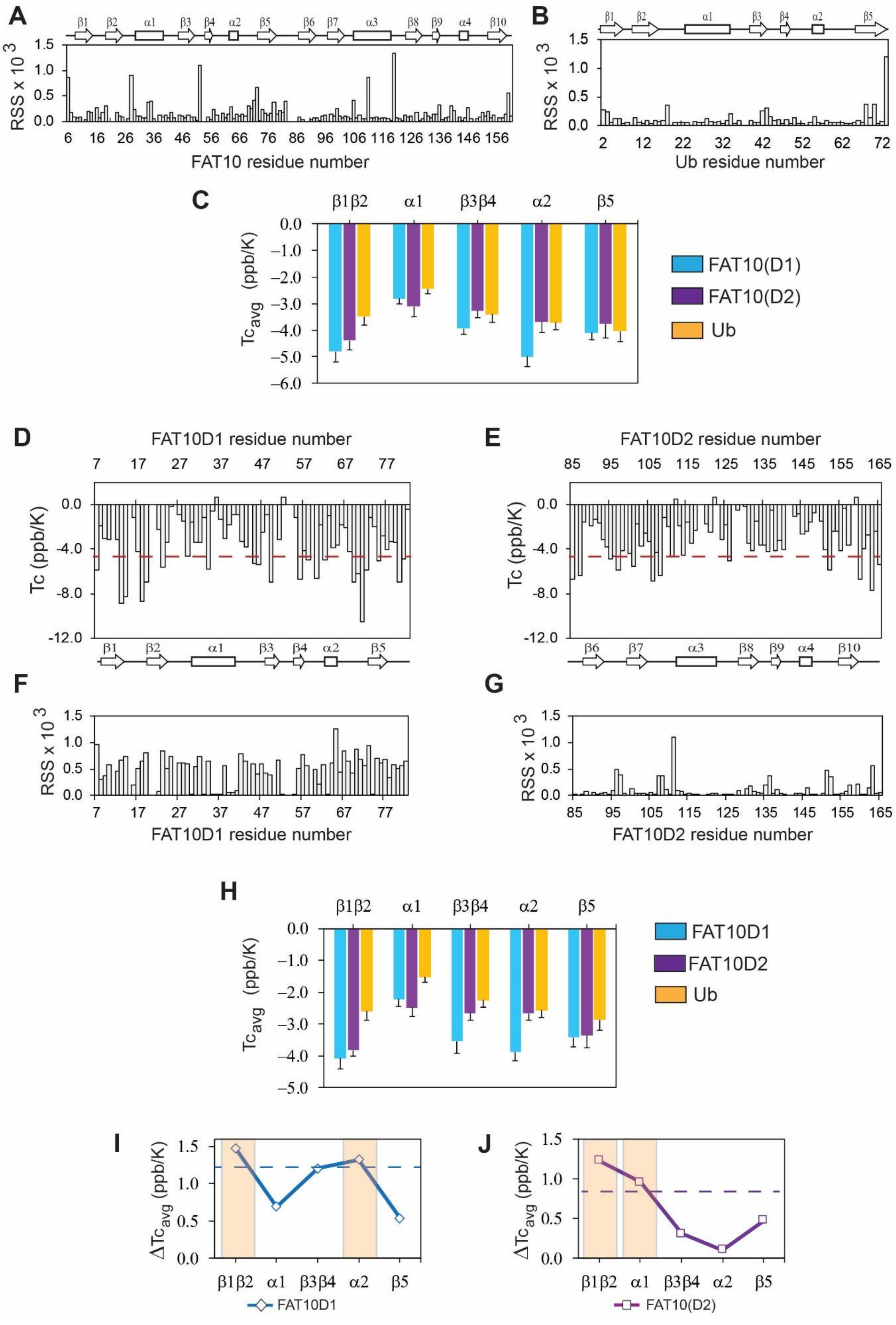

A
B
C
D
E
F
G
H
I
J
RSS x 10 3
FAT10 residue number
Ub residue number
Tcavg (ppb/K)
FAT10(D1)
FAT10(D2)
Ub
FAT10D1 residue number
FAT10D2 residue number
Tc (ppb/K)
FAT10D1
FAT10D2
ΔTcavg (ppb/K)
FAT10(D2)

Figure S11. The temperature dependence of amide chemical shifts. (A) The Residual Sum Square (RSS) for each residue of Fat10 and (B) ubiquitin were calculated and plotted. (C) The averaged Tc of Fat10 and ubiquitin over various protein segments. Fat10(D1) and Fat10(D2) are two domains in Fat10. The ubiquitin segments are defined as β1β2: 2-22; α1: 23-34; β3β4: 35-50; α2: 51-63, and β5: 64-74. For the D1 domain in full-length Fat10, Fat10(D1), β1β2: 8-28, α2: 29-42, β3β4: 43-56, α2: 57-72 and β5: 73-85. For the D2-domain in full-length Fat10, Fat10(D2), β1β2: 85-110, α2: 111-121, β3β4: 122-143, α2: 144-153 and β5: 154-160. (D) and (E) are the Tc values of isolated Fat10 domains. The broken red lines denote Mean+SD. (F) and (G) errors for Tc calculation for the isolated Fat10 domains. (H) The averaged Tc of Fat10 domains and ubiquitin over various protein segments. Ub values are replotted here for comparison. (I) The difference in averaged temperature coefficients ($\Delta Tc_{avg}$) between the isolated N-terminal domain and Ub, where $\Delta Tc_{avg} = Tc(Ub)_{avg} - Tc(Fat10D1)_{avg}$. The blue line is mean + error. (D) The difference in averaged temperature coefficients ($\Delta Tc_{avg}$) between the isolated C-terminal domain and Ub, where $\Delta Tcavg = Tc(Ub)_{avg} - Tc(Fat10D2)_{avg}$. The purple line is the mean + error.

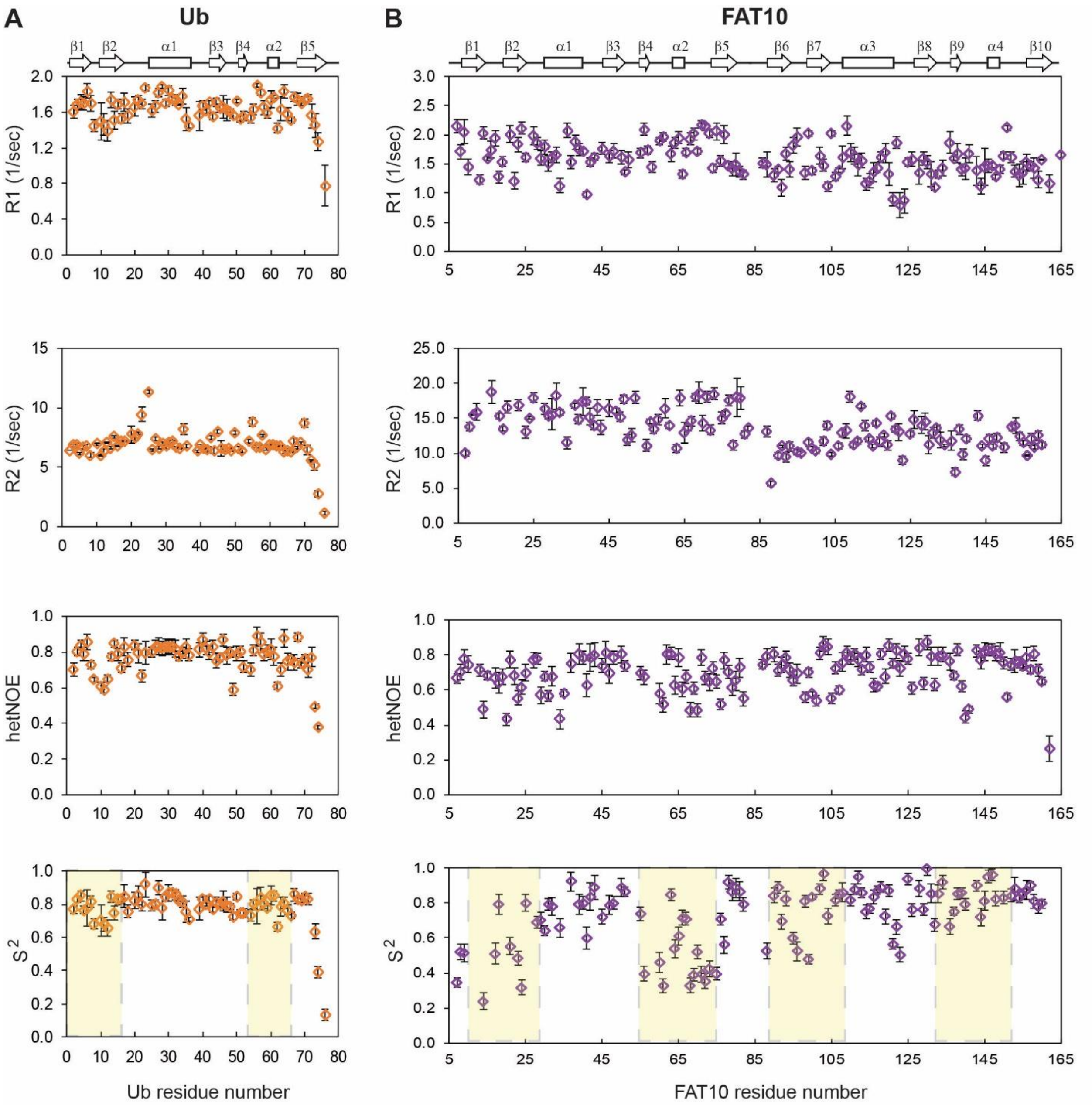


Figure S12. Measurement of backbone dynamics in Fat10 and Ub. (A) The measured values of longitudinal relaxation rates (R1), transverse relaxation rates (R2), Nuclear Overhauser Enhancement (hetNOE), and order parameter ($S^2$) are plotted for Ub. The ubiquitin secondary structure elements are provided on the top panel. (B) The measured R1, R2, hetNOE, and S2 values are plotted for the Fat10. The Fat10 secondary structure elements are provided on the top panel. The measured values are collected at the field of 800 MHz.

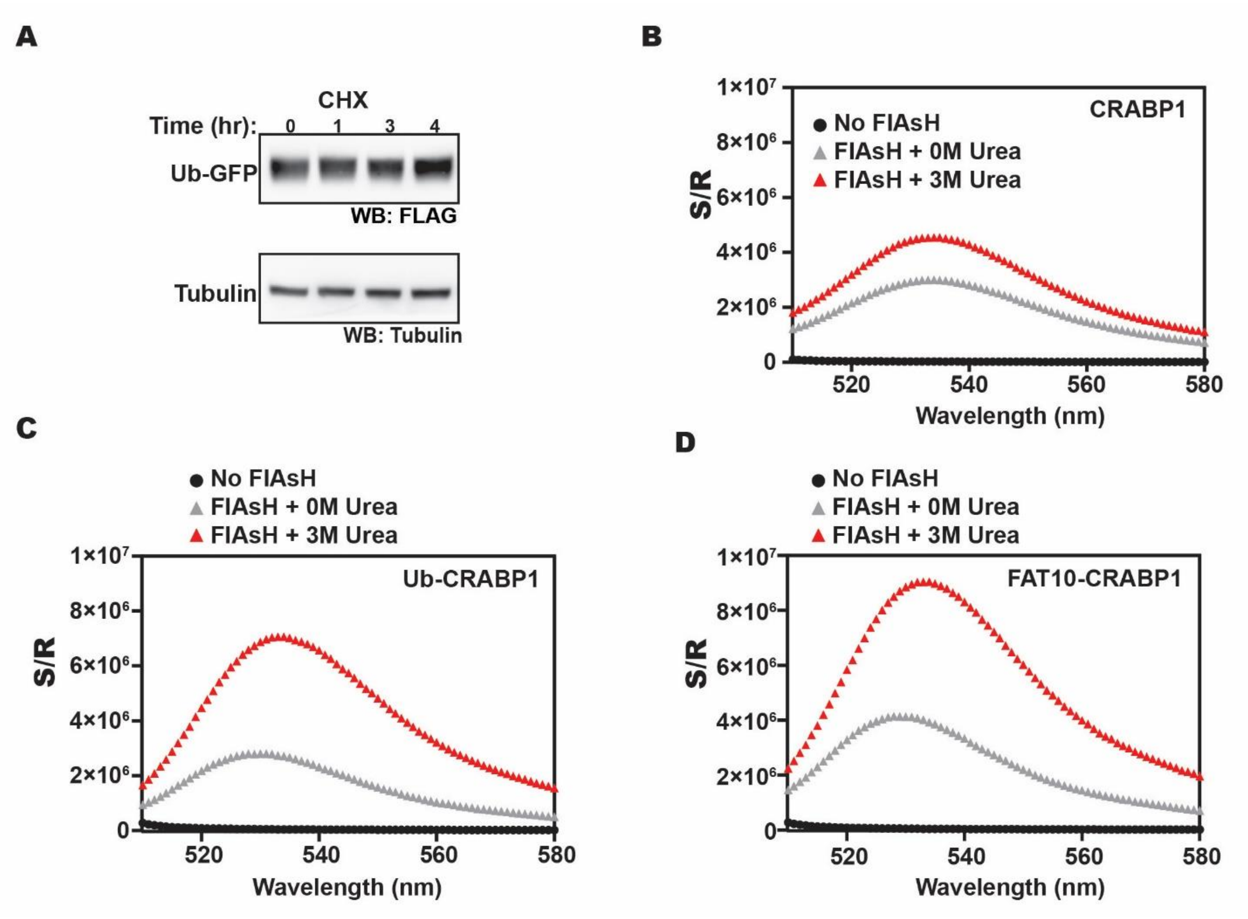


Figure S13. (A) In-vivo degradation of Ub-GFP after cycloheximide treatment over different time points. Tubulin is used as the loading control. (B) *The in-vivo* fluorescence signal of free-CRABP1, (C) Ub-CRABP1, and (D) Fat10-CRABP1 when cells were incubated without or with 3M urea.

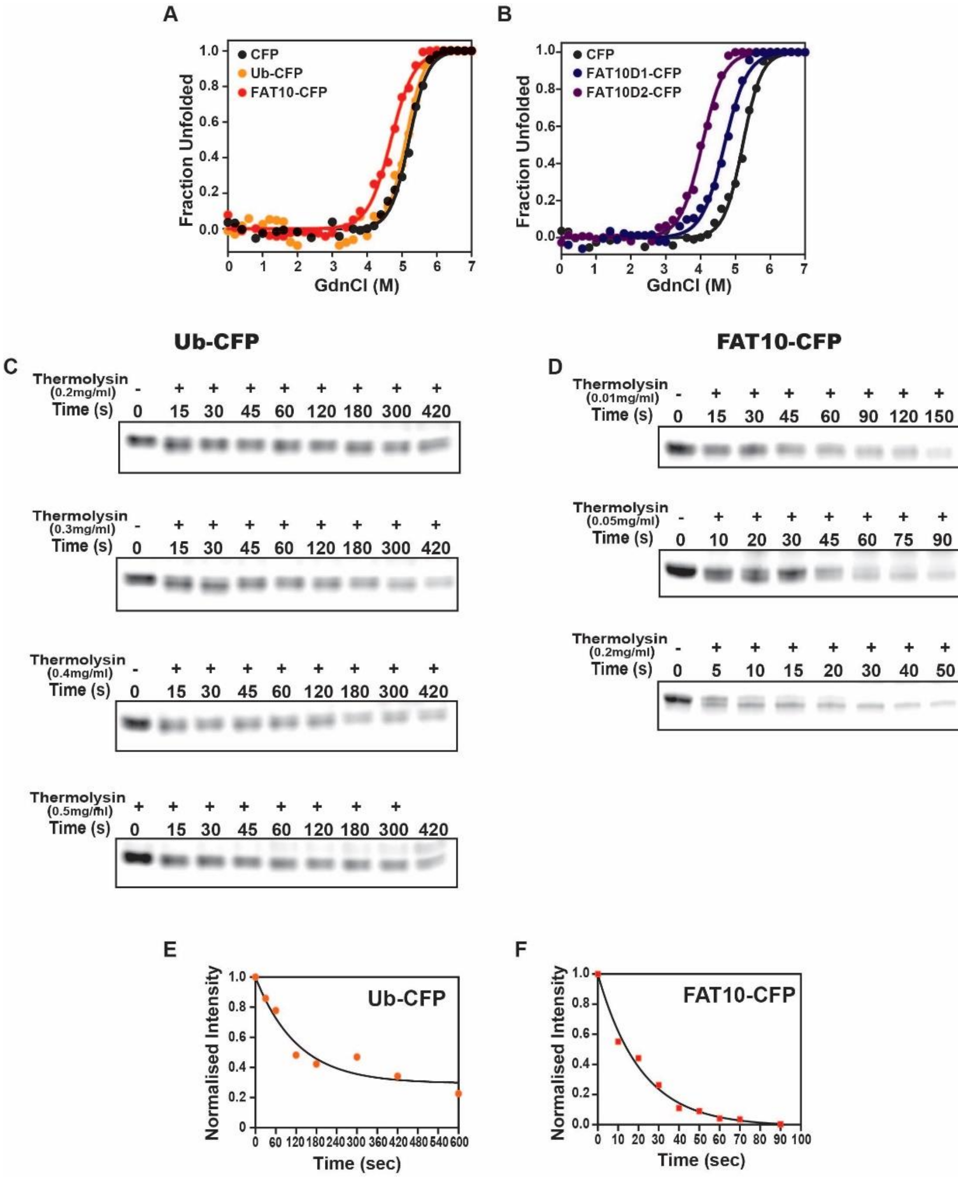


Figure S14. (A) GdnCl melt curves of CFP, Ub-CFP, and Fat10-CFP where only CFP was probed using its fluorescence signal. The fluorescence values were normalized and plotted against GdnCl concentration. (B) Similarly, the GdnCl melt curve for Fat10D1-CFP and Fat10D2-CFP was plotted. The CFP melt curve in (A) is repeated here for comparison. (C) In-gel fluorescence image for native-state proteolysis of Ub-CFP treated with 0.2mg $ml^{-1}$, 0.3mg $ml^{-1}$, 0.4mg $ml^{-1}$and 0.5mg $ml^{-1}$ of Thermolysin. (D) In-gel fluorescence image for native-state proteolysis of Fat10-CFP treated with 0.01mg $ml^{-1}$, 0.05mg $ml^{-1}$, and 0.2mg $ml^{-1}$ of Thermolysin. The normalized intensity of in-gel fluorescence of CFP for 0.1 mg/ml thermolysin treatment is plotted against time for (E) Ub-CFP and (F) Fat10-CFP.

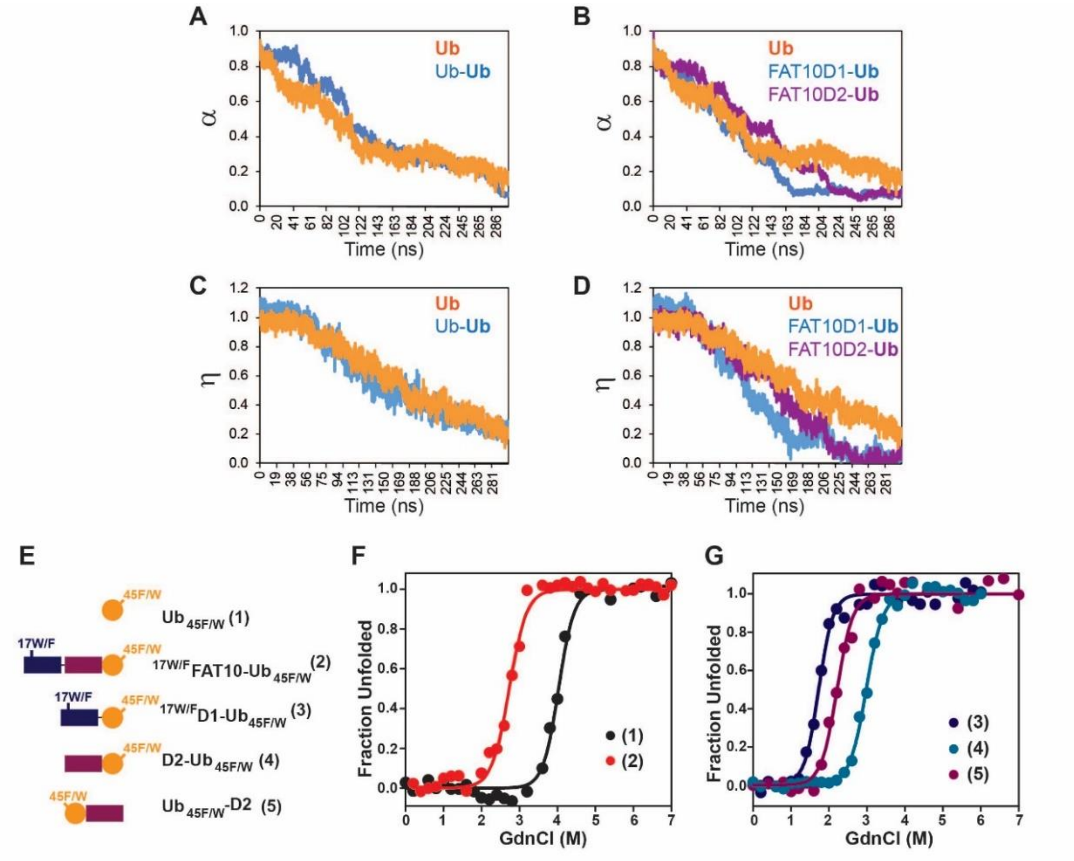


**H**

| | Ub | FAT10-Ub | FAT10D1-Ub | FAT10D2-Ub | Ub-FAT10D2 |
|---|---|---|---|---|---|
| $C_m$ (M GdnCl) | 4.03 ± 0.02 | 2.74 ± 0.03 | 1.73 ± 0.02 | 2.98 ± 0.02 | 2.21 ± 0.02 |
| $\Delta C_m$ (M GdnCl) | | 1.29 ± 0.04 | 2.30 ± 0.03 | 1.05 ± 0.03 | 1.82 ± 0.03 |
| m (kcal $mol^{-1}$ $M^{-1}$) | 2.89 ± 0.38 | 2.57 ± 0.21 | 3.13 ± 0.27 | 2.68 ± 0.14 | 2.82 ± 0.30 |
| Δm (kcal $mol^{-1}$ $M^{-1}$) | | -0.32 ± 0.43 | -0.24 ± 0.47 | -0.21 ± 0.41 | -0.07 ± 0.48 |
| $\Delta G_{unfolding}$ (kcal $mol^{-1}$) | 11.68 ± 1.52 | 7.03 ± 0.58 | 5.40 ± 0.47 | 8.01 ± 0.42 | 6.24 ± 0.67 |
| $\Delta\Delta G_{unfolding}$ (kcal $mol^{-1}$) | | -4.68 ± 1.63 | -6.28 ± 1.59 | -3.67 + 1.58 | -5.44 + 1.66 |

Figure S15. (A) Simulations of Mono-ubiquitin (Ub) and di-ubiquitin ($Ub_2$), Fat10D1-Ub, and Fat10D2-Ub were performed at 450K. The fraction of native contacts is defined as (α) and plotted against the time for ubiquitin and the proximal unit in di-ubiquitin $Ub_2$ (D2). (B) α is plotted for ubiquitin in the Fat10D1-Ub, Ub in Fat10D2-Ub, and monoUb. (C) The fraction of native backbone hydrogen bonds in the beta-sheet β1 to β5 is defined as (η) and plotted against the time for ubiquitin and the proximal unit in di-ubiquitin $Ub_2$ (D2). (D) is the same as (C), plotted for ubiquitin in the Fat10D1-Ub, ubiquitin in Fat10D2-Ub, and monoUb. (E) ubiquitin varieties used in this study where Ub45F/W are covalently conjugated at the C-terminus of $_{17W/F}$Fat10 and $_{17W/F}$Fat10D1 and Fat10D2. $Ub_{45F/W}$ was also fused at the N-terminus of Fat10D2. (F) GdnCl melt curves of $Ub_{45F/W}$ and $_{17W/F}$Fat10-$Ub_{45F/W}$ are shown. The tryptophan fluorescence signals of $Ub_{45F/W}$ were normalized and plotted against the GdnCl concentration. (G) GdnCl melt curves were plotted for $_{17W/F}$Fat10D1-$Ub_{45F/W}$, Fat10D2-$Ub_{45F/W,}$ and $_{45F/W}$Ub-Fat10D2. (H) A table with details of thermodynamic parameters of $Ub_{45F/W}$ in free and bound forms is shown.

**A**

| | Number of Intermolecular Contacts between FAT10 and substrate ubiquitin detected per μsec |
|---|---|
| FAT10-Ub | 4140258 |
| FAT10D2-Ub | 4441543 |
| Ub-FAT10D2 | 6355889 |

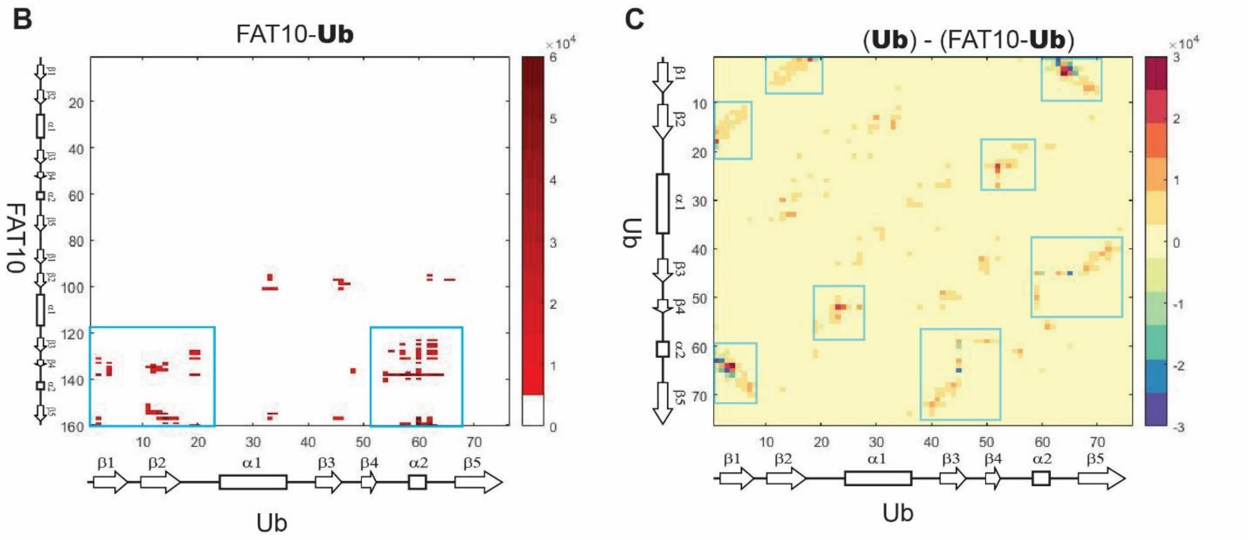


Figure S16. (A) The number of intermolecular contacts (within 4.5 Å) per μsec between Fat10 and substrate ubiquitin in the Fat10-Ub conjugate detected in the MD simulation. The same is calculated for the conjugate of the D2 domain and ubiquitin in the Fat10D2-Ub and Ub-Fat10D2 conjugates. (B) The intermolecular contacts between Fat10 and ubiquitin are plotted as a contact map. Blue rectangles mark the contact with significant occupancies. (C) The difference between long-range (i, i+5) intra-ubiquitin contacts in free ubiquitin and Fat10-ubiquitin conjugate is plotted as a contact map. Positive values indicate ubiquitin contacts disrupted in the Fat10-Ub conjugate, and negative values indicate new ubiquitin contacts formed in the Fat10-Ub conjugate.

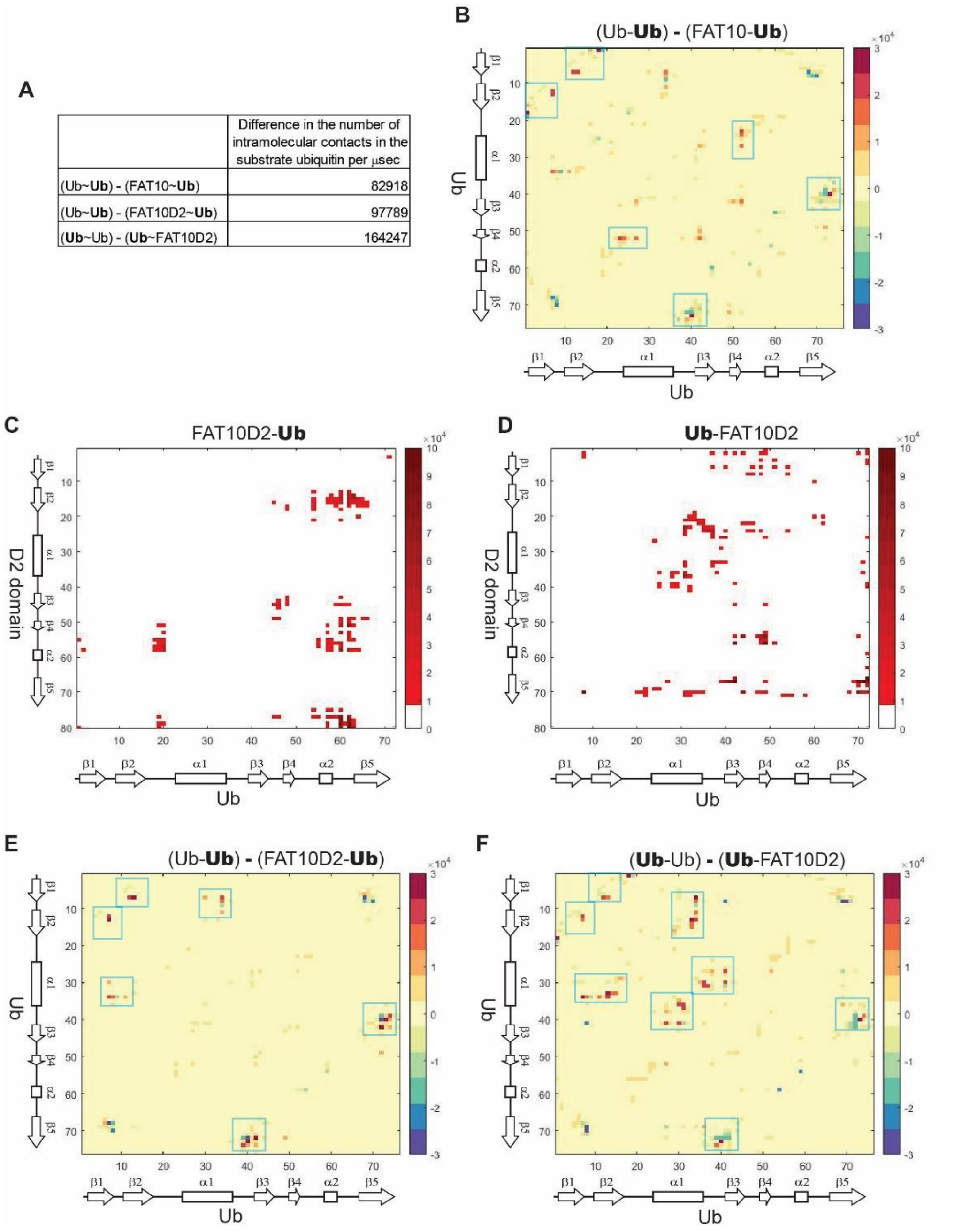


Figure S17. (A) The difference of long-range (i, i+5) intra-ubiquitin contacts in diubiquitin and Fat10-ubiquitin conjugates are given as a table. The positive value of difference indicates contacts present in diubiquitin but disrupted in Fat10-ubiquitin. (B) The difference between long-range intra-ubiquitin contacts in diubiquitin and Fat10-ubiquitin conjugate is plotted as a ubiquitin contact map. Positive values indicate contacts disrupted in the Fat10-ubiquitin conjugate but present in diubiquitin. Negative values indicate new contacts formed in Fat10-ubiquitin but absent in diubiquitin. Blue rectangles indicate regions with severely disrupted contacts in Fat10-ubiquitin. (C) and (D) The intermolecular contacts are plotted between ubiquitin and D2 domain for Fat10D2-Ub and Ub-Fat10D2, respectively. (E) and (F) The difference of intra-ubiquitin contacts is plotted for Fat10D2-ubiquitin and ubiquitin-Fat10D2 in, respectively. Positive values indicate contacts disrupted in the Fat10-ubiquitin conjugates but present in diubiquitin. Negative values indicate the inverse. Blue rectangles indicate regions with severely disrupted contacts in the Fat10D2 conjugates.

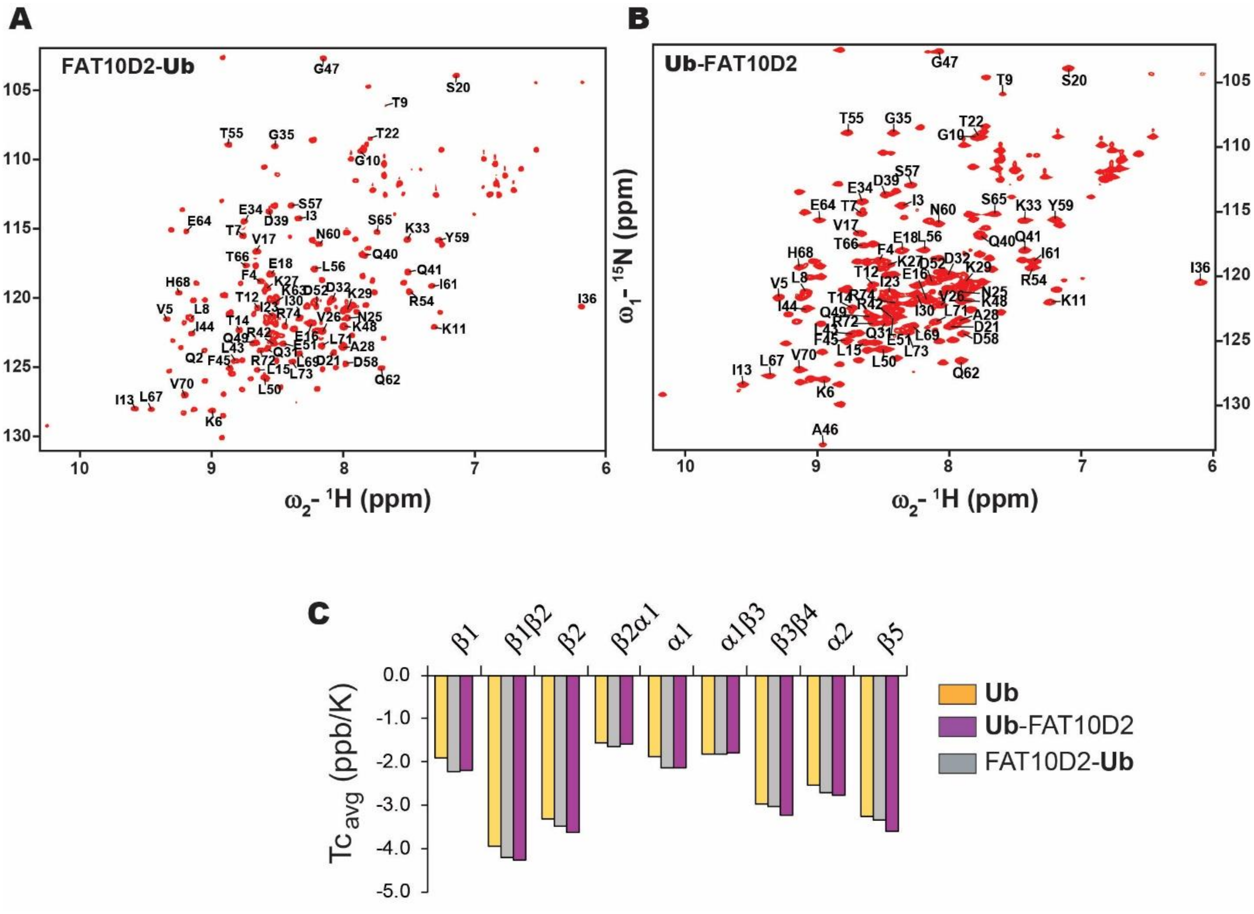


Figure S18. 2D-HSQC spectrum with assigned peaks of ubiquitin residues in each of the chimeric constructs: (A) Fat10D2-Ub and (B) Ub-Fat10D2. (C) The averaged Tc is plotted for the various regions in ubiquitin for the free ubiquitin, and ubiquitin conjugated to Fat10 domains.

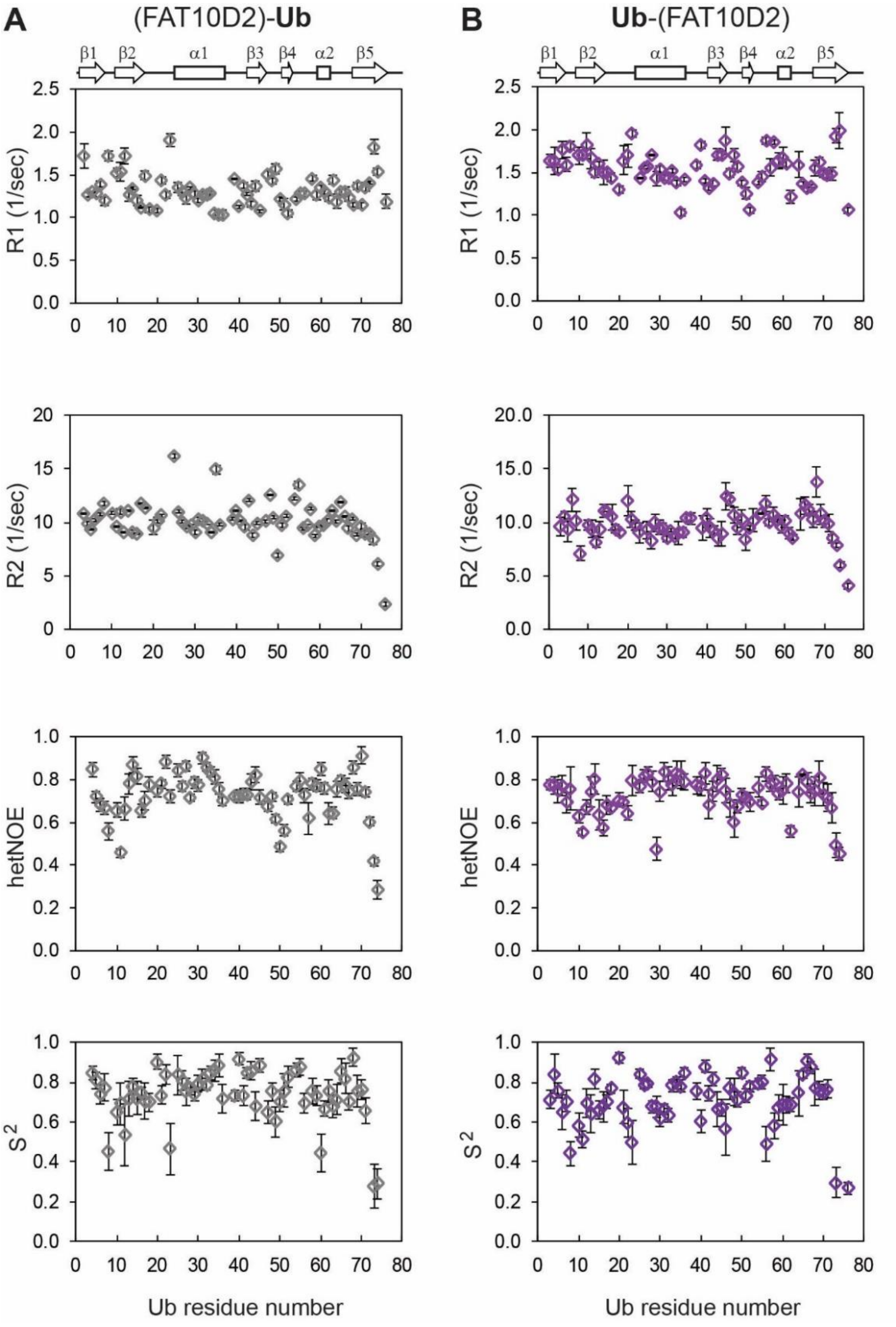


Figure S19. Changes in the ubiquitin backbone dynamics by conjugation to Fat10 domains. The measured values of longitudinal relaxation rates (R1), transverse relaxation rates (R2), Nuclear Overhauser Enhancement (hetNOE), and order parameter ($S^2$) for ubiquitin residues are shown in the case of free (A) Fat10D2-Ub and (B) Ub-Fat10D2. The Ub secondary structure elements are provided on the top panel.

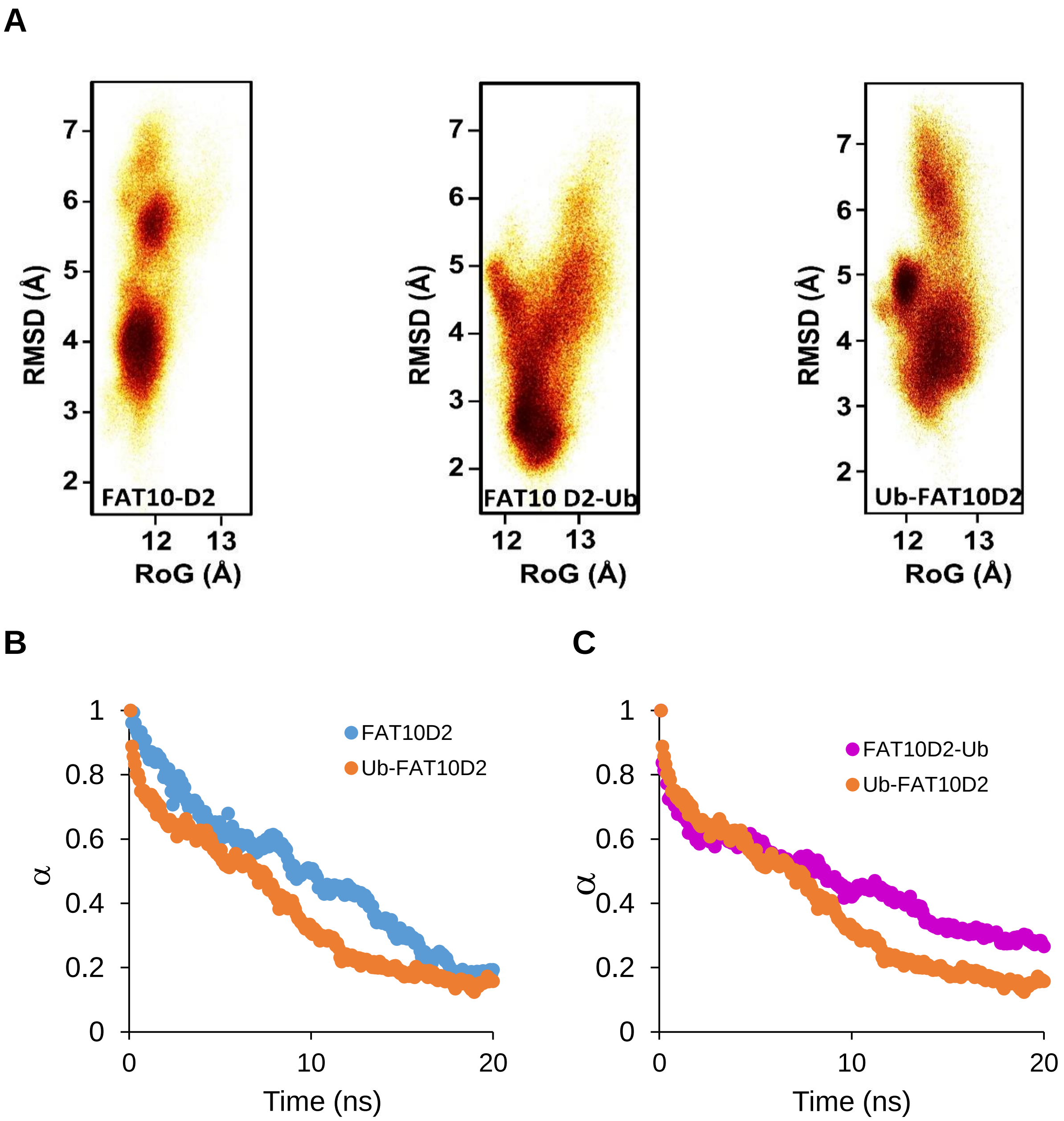


Figure S20. (A) The free energy landscape of Fat10D2 in the free form, Fat10D2-Ub, and Ub-Fat10D2 is plotted as a function of RMSD and radius of gyration (Rgyr), which was obtained from simulations across three replicas (3 x 2.5 μs) performed at 300K. (B) Simulations of free Fat10D2 and Ub-Fat10D2 were performed at 450K. The fraction of native contacts is defined as (α) and plotted against time. The data is averaged over ten replicas. (C) Similar to (B) plotted for Ub-Fat10D2 and Fat10D2-Ub. The Ub-Fat10D2 data are replotted here as a control.

**Legends for movies**

**Movie S1.** Mechanical unfolding of Fat10 and di-ubiquitin by MD simulations is shown here. The top panel is Fat10, and the bottom panel is di-ubiquitin. The PMF values of the two molecules are provided in between.

**Movie S2.** The thermal unfolding of Fat10 domains and ubiquitin is shown here. Fat10D1 is on the left and colored blue. Fat10D2 is colored purple and displayed in the middle. Ubiquitin is shown at the right and colored orange. The top panel is Fat10, and the bottom panel is di-ubiquitin. The PMF values of the two molecules are provided in between. The duration of simulations is shown at the bottom right. The kinetics of unfolding Fat10 domains are significantly faster than Ub.
